# Co-translational assembly counteracts promiscuous interactions

**DOI:** 10.1101/2021.07.13.452229

**Authors:** Maximilian Seidel, Anja Becker, Filipa Pereira, Jonathan J. M. Landry, Nayara Trevisan Doimo de Azevedo, Claudia M. Fusco, Eva Kaindl, Janina Baumbach, Julian D. Langer, Erin M. Schuman, Kiran Raosaheb Patil, Gerhard Hummer, Vladimir Benes, Martin Beck

## Abstract

During the co-translational assembly of protein complexes, a fully synthesized subunit engages with the nascent chain of a newly synthesized interaction partner. Such events are thought to contribute to productive assembly, but their exact physiological relevance remains underexplored. Here, we examined structural motifs contained in nucleoporins for their potential to facilitate co-translational assembly. We experimentally tested candidate structural motifs and identified several previously unknown co-translational interactions. We demonstrate by selective ribosome profiling that domain invasion motifs of beta-propellers, coiled-coils, and short linear motifs act as co-translational assembly domains. Such motifs are often contained in proteins that are members of multiple complexes (moonlighters) and engage with closely related paralogs. Surprisingly, moonlighters and paralogs assembled co-translationally in only one but not all of the relevant assembly pathways. Our results highlight the regulatory complexity of assembly pathways.

## Introduction

Protein complexes are a key organizational unit of the proteome. Their modular composition has facilitated the evolution of a very diverse repertoire of folds and corresponding functions. To maintain this very diverse repertoire within the crowded cellular environment poses a logistic burden as the energy gap favoring specific over non-specific binding decreases with proteome complexity (Johnson and Hummer, 2011). Therefore, it has been proposed that assembly pathways impose a major restraint on the evolution of protein complexes (Marsh et al., 2013), whereby duplication events of subunits during divergent evolution may necessitate the diversification of protein interfaces or sophisticated quality control mechanisms to avoid promiscuous binding (Mena et al., 2020). Co-translational interactions of nascent polypeptides with their respective binding partner have been discovered for many eukaryotic protein complexes (Bertolini et al., 2021; Duncan and Mata, 2011; Kamenova et al., 2019; Shiber et al., 2018). Homomeric complexes may assemble by co-co assembly in which either nascent chains emerge from consecutive ribosomes of the same mRNA entangled in *cis*, or alternatively from multiple mRNAs that are clustered by nascent chain interactions in *trans* (Bertolini et al., 2021; Natan et al., 2018). In contrast, many heteromers may rely on co-post assembly (Duncan and Mata, 2011; Kamenova et al., 2019; Kramer et al., 2019) which we will further refer to as co-translational assembly. Here, a soluble, fully synthesized subunit binds to the nascent polypeptide chain of the interactor. Such co-translational assembly events contribute to orphan protein stability and solubility and may be coordinated with the association of assembly chaperones (Shiber et al., 2018). It has been proposed that they may be beneficial for nascent chain folding or non-promiscuous stoichiometric assembly (Kramer et al., 2019) and have been hypothesized to seed assembly pathways when moonlighting interactions are possible (Schwarz and Beck, 2019). Since co-translational assembly pathways of moonlighters remain largely unexplored, the exact physiological contribution of distinguished co- and/or post-translational assembly pathways remains uncertain.

Nuclear pore complexes (NPC) perforate the nuclear envelope (NE) to facilitate nucleocytoplasmic exchange. They are among the largest, non-polymeric, eukaryotic assemblies and are composed of ^~^30 different nucleoporins (Nups) that constitute a multi-layered modular architecture of astonishing complexity (Beck and Hurt, 2017; Lin and Hoelz, 2019). Beyond canonical protein interfaces of nucleoporin subcomplexes of up to 10 components, various other types of interactions are crucial for the formation of the higher ordered, 8-fold rotational symmetric structure of ^~^500 nucleoporins in yeast (Hampoelz et al., 2019a; Lin and Hoelz, 2019). Those include weak interactions of intrinsically disordered Phenylalanine-Glycine (FG)-rich repeats contained in so-called FG-Nups that function as a velcro (Onischenko et al., 2017) and short linear motifs (SliMs) within so-called linker Nups that facilitate interactions within and across subcomplexes (Fischer et al., 2015; Stuwe et al., 2015a). Furthermore, structured motifs such as beta-propeller complementation (Brohawn and Schwartz, 2009; Debler et al., 2008; Nagy et al., 2009) and coiled-coil interactions (Chug et al., 2015; Kim et al., 2018; Stuwe et al., 2015a) are observed in multiple instances. Interestingly, different Nup subcomplexes that have evolved from each other may contain shared or closely related subunits that assemble promiscuously *in vitro* (Bailer et al., 2001; Melčák et al., 2007; Solmaz et al., 2011; Ulrich et al., 2014). Nevertheless, it remains unclear how such promiscuous interactions are suppressed or discriminated *in vivo*.

Due to the importance of the NPC as a permeability barrier, faithful assembly throughout the cell cycle imposes a challenge for cells which is addressed by different pathways depending on the spatiotemporal context (Hampoelz et al., 2019a). While NPCs are made from pre-existing building blocks during post-mitotic assembly in higher eukaryotes, they are synthesized from scratch during the ubiquitous interphase assembly pathway (Hampoelz et al., 2019a) and *Drosophila* oogenesis (Hampoelz et al., 2016, 2019b). Interphase assembly is the only known biogenesis pathway in yeast and spatially proceeds from the inside-out at the NE (Otsuka and Ellenberg, 2018). Although, the rough order of subcomplex recruitment to membranes has been resolved (Hakhverdyan et al., 2021; Hampoelz et al., 2019a; Onischenko et al., 2020; Otsuka and Ellenberg, 2018), little is known about the early steps of assembly that may occur away and independently from membranes. Besides local (Hampoelz et al., 2019b) and some co-translational events (Lautier et al., 2021) that have been discovered during NPC assembly, it remains unclear which of the above-introduced motifs are subject to such events, in which order they intertwine into the assembly pathways, and how exactly they contribute to faithful assembly. Here, we have elucidated the role of a subset of such motifs for the co-translational *de novo* assembly of NPC subcomplexes during interphase assembly in *Saccharomyces cerevisiae*.

## Results

The competition between the specific interactions stabilizing a complex and the far more numerous non-specific promiscuous interactions intensifies with increasing numbers of complex subunits and possible interaction partners (Johnson and Hummer, 2011). Here, we extend this theory by including co-translational assembly as a possible assembly enhancer. The assembly pathways of higher-ordered structures, such as a hypothetical complex ABCD built from the protein components A, B, C, and D (**Figure 1**), have to overcome two predominant obstacles: (i) intermediates such as AB and ABC may fall apart prior to further stabilization by the subsequent binding partner; and (ii) each component will compete with non-native interactors X for binding. If such aggregates with non-native interactors indeed removed a considerable fraction of assembly components and intermediates, the assembly would become ineffective. Co-translational assembly may resolve or at least ameliorate assembly errors by generating a sequential assembly pathway in which components arrive one after the other. In a simplified model, we propose that the yield of successful assembly depends on the competition between assembly and aggregation in which the latter is driven by the incorrect folding or inclusion of non-native interactors X (**Figure 1**). Hence, the overall yield decreases exponentially as (1-*p*)^*N*-1^ where *p* expresses the probability of incorrect incorporation and *N* the number of protein components. Notably, X may account for off-pathway intermediates with randomly colliding proteins but also for competition with paralogs.

**Figure 1:**
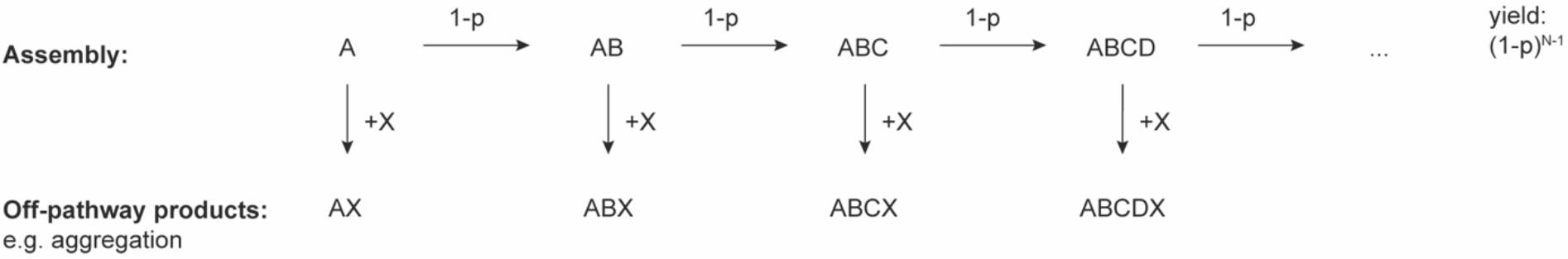
Theoretical benefits of co-translational assembly. In a linear schematic of co-translational assembly, we assume that the probability for incorrect incorporation at each step is *p*, while correct assembly occurs with a probability of 1-p. We further assume that incorrect incorporations of X lead to the removal of material from the assembly process. The yield of assembly thus decreases exponentially with the number of components *N*.

In this model, a fundamental challenge is that the probability of correct assembly with B, C, D, … must be much higher than the one for incorrect assembly with X, 1 − *p* ≫ *p*, even though a vast amount of incorrect binders X may have at least some binding affinity. Co-translational assembly can tolerate higher levels of such competitive binders because it increases the dwell-time of synthesis intermediates at the ribosome in which non-binding surfaces are not yet exposed. Most importantly, the larger the number of subunits *N*, the smaller *p* must be to ensure high yields. Therefore, the more subunits a given complex has, the less likely it is to assemble post-translationally at a high yield. This imposes a severe challenge for the biogenesis of very large complexes with many subunits such as the NPC (**Figure 2a**) and is further complicated by the fact that parts of the biogenesis pathway have to proceed to the nucleus in the absence of translating ribosomes. One would thus predict that cells harness the power of co-translational assembly for the biosynthesis of NPC subcomplexes in the cytosol.

**Figure 2:**
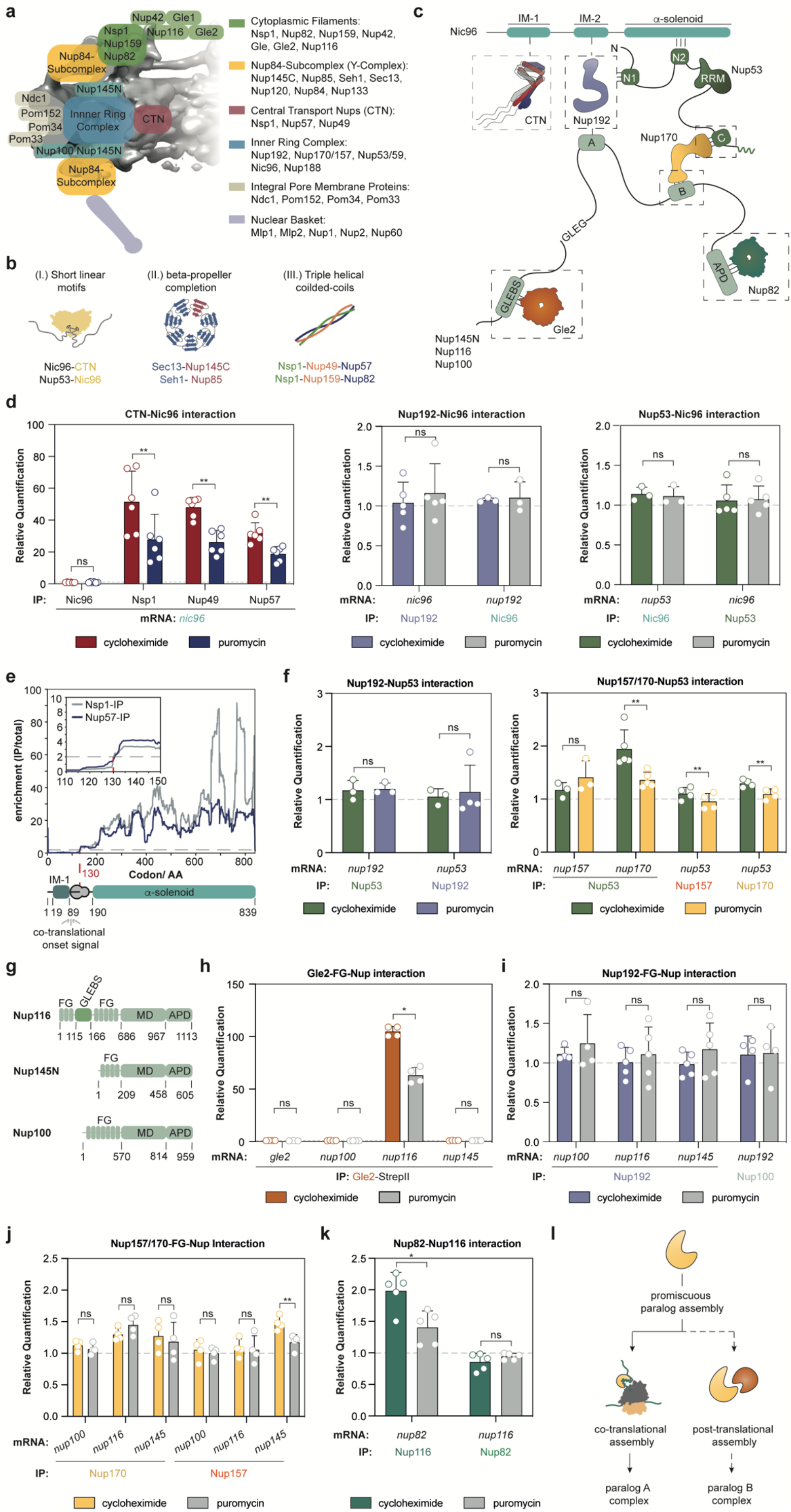
Co-translational assembly of linker Nups. **a**, Scheme of the NPC architecture. Subcomplexes are mapped to the *in cellulo* structure of the *S. cerevisiae* NPC (EMD: 10198) (Allegretti et al., 2020). **b**, Recurring structural features of the NPC: (i) high density of short linear motifs and small structured domains, (ii) incomplete beta-propellers in the Nup84-subcomplex and (iii) triple helical coiled-coils. **c**, Scheme of previously determined interaction motifs contained in linker Nups that may help to organize the assembly *in vivo*. Figure adapted from Beck and Hurt (2017). **d**, RIP-qPCR experiments with affinity purifications of CTN/Nic96 (left) that associate via the IM-1, Nup192/Nic96 (middle) that interact via the IM-2 and Nup53/Nic96 that interact via N2 (right), imply that the entire CTN binds co-translationally to Nic96. **e**, SeRP experiments with affinity purifications of Nsp1 and Nup57 reveal a synchronous co-translational binding within the *nic96*-transcript. Data were derived from four individual biological replicates. **f**, RIP-qPCR for Nup192 and Nup53 (left) that interact via the N1 motif and for Nup157/170 and Nup53 (right) that interact using the C-motif. **g**, Primary structure scheme of the paralogous FG-Nups Nup145N, Nup116 and Nup100. **h**, RIP-qPCR experiments with affinity purified Gle2 that show co-translational binding to nascent Nup116. RIP-qPCR was used to characterize co-translational interactions between **i**, Nup192 and Nup100 that bind to each other by the A-motif, **j**, Nup157 and Nup170 that could provide the B-motif for binding and **k**, Nup116 and Nup82 that interact via the APD. **l**, Scheme highlighting that designated assembly pathways may exist for paralogs. Bar graphs in panel **d, f**, and **h-k** depict mean ± SD from three to six biologically independent experiments. ns p>0.05,* p<0.05, ** p<0.01 (Two-sided, paired t-test). AA: amino acid, IP: immunoprecipitation.

We surveyed the known structural repertoire of nucleoporins for domains that could potentially engage in co-translational interactions because they (i) are small interaction motifs found in linker Nups; (ii) have obviously complemented the fold of another nucleoporin; or (iii) are shared between multiple complexes and thus could be promiscuous interactors (**Figure 2b**). This concerns either the competitive binding of Nups for the same binding domains within the NPC or the integration of Nups into distant, functionally unrelated complexes. In order to elucidate the implementation of co-translation for faithful assembly of the NPC, we experimentally validated these motifs in a hypothesis-driven approach.

To analyze co-translational interactions, we first generated a library of C-terminally Twin-StrepII tagged (Schmidt et al., 2013) Nups using a scar-free cloning technique in *S. cerevisiae* (Carvalho et al., 2013). Scar-free cloning preserves the endogenous 3’ untranslated region (3’UTR) of a messenger RNA and avoids changes that may affect mRNA fate and translation (Mayr, 2017). We used these strains for affinity purification of the respective StrepII-tagged bait (**Figure S1a**) and analyzed the co-enriched mRNAs by quantitative real-time PCR adapting previously established methods (Duncan and Mata, 2011; Kamenova et al., 2019; Shiber et al., 2018). Below we refer to this method as RIP-qPCR (**Figure S1b**). As a positive control, we reproduced the known co-translational interaction of fatty acid synthase subunit Fas1 with nascent Fas2 (Shiber et al., 2018), represented by enrichment of *fas2*-mRNA. This interaction was sensitive to the disruption of translation by puromycin, which causes dissociation of the nascent chain (Blobel and Sabatini, 1971), and inverse tagging of Fas2 instead of Fas1 did not enrich for either mRNA, as expected (**Figure S1c-S1e**).

Using the above standards for the experimental validation, we tested the potential co-translational interactions of various full length Nups that engage with linker Nups. First, we investigated the co-translational landscape of Nic96 which contains several motifs to connect the inner ring to the nuclear envelope and the central transport Nup (CTN) trimer (**Figure 2c**) (Fischer et al., 2015; Lin et al., 2016; Stuwe et al., 2015a). Our RIP-qPCR data revealed a novel and strong co-translational interaction of Nsp1, Nup49, and Nup57 with the nascent chain of Nic96 (**Figure 2d**) suggesting that the fully assembled CTN trimer is recruited at once to nascent Nic96. Inverse mRNA enrichment for any subunit of the CTN in Nic96 RIP-qPCR experiments was not observed (**Figure S2a**). Despite co-translational interactions at the predicted IM-1 motif of Nic96, we did not detect any other co-translational events with Nup192 or Nup53 (**Figure 2d**).

To illustrate that the IM-1 domain is sufficient to recruit Nsp1 and Nup57 to nascent Nic96, we turned to selective ribosome profiling (SeRP), as previously described (Shiber et al., 2018) (**Figure S3 and S4a**). This method relies on the comparison of ribosome-protected mRNA footprints from a total translatome to those that are affinity-purified using a specific bait protein that may engage in interactions with nascent chains. Thereby the onset of the co-translational interaction within a given open reading frame (ORF) is revealed. As a positive control, we reproduced the co-translational onset of the interaction of full length Fas1 with the nascent chain of Fas2. We observed a very strong enrichment of footprints within the *fas2*-transcript precisely at the previously reported onset (Shiber et al., 2018) at the Fas2 assembly domain (**Figure S4b-S4c**).

The selective ribosome profiles generated from Nsp1- and Nup57-IPs for *nic96*-mRNA show simultaneous binding properties to the nascent chain at codon 130. Considering a 30 to 40 amino acid delay of nascent chain accessibility due to the ribosome exit tunnel (Fedyukina and Cavagnero, 2011), the measured onset coincides with the exposure of the IM-1 motif of Nic96 from the exit tunnel (**Figure 2e**). The synchronous onset further underscores the notion that only fully assembled CTN trimer can associate with Nic96, in line with previous biochemical analysis (Chug et al., 2015; Fischer et al., 2015; Schlaich et al., 1997). SeRP of Nic96-IPs further indicated that Nic96 does not engage with nascent chains of the other known interactors (**Figure S5a**).

Independent of Nic96, Nup53 and Nup192 can also directly associate with one another via an N1 interaction site of Nup53 (Stuwe et al., 2014) (**Figure 2c**). Our targeted RIP-qPCR approach revealed that this interaction occurs independently of co-translational pathways (**Figure 2f**). In contrast, Nup53 that binds to Nup170/157 with the so-called C-motif (Lin et al., 2016) co-translationally binds to nascent Nup170 but not Nup157 (**Figure 2f**). Although Nup170 and Nup157 are paralogs, the assembly pathways with Nup53 are divergent. Taking into account biochemical data (Onischenko et al., 2009), it appears plausible that Nup53 evolved designated assembly pathways for both Nup157 and Nup170 to compensate for the risk of unwanted subcomplexes. Nup157 and Nup170 also generated a puromycin-sensitive signal for *nup53*-mRNA but with very small effect size.

We next wanted to understand whether the paralogous linker proteins, Nup100, Nup116, and Nup145N are involved in co-translational subcomplex formation. The N-terminal GLEBS domain of Nup116 binds to Gle2 and is absent in Nup100 or Nup145N (Bailer et al., 1998) (**Figure 2g**). We found that Gle2 binds co-translationally to the nascent chain of Nup116 but not Nup145N or Nup100 (**Figure 2h**) and reasoned that the GLEBS domain arose to specify interactions of Nup116. Next, we analyzed interactions of the three above-introduced linker Nups with Nup192 and Nup170/157 that are mediated by the A-motif and B-motif further downstream (Lin et al., 2016) (**Figure 2c**). We did not detect any co-translational interactions with Nup192 (**Figure 2i**) and only weak co-translational interactions of Nup157 with nascent Nup145N (**Figure 2j**). This finding is reminiscent of the assembly pathways of Nup53 with Nup170/157. While Nup145N can interact with both, Nup170 and Nup157 (Fischer et al., 2015; Lutzmann et al., 2005), the assembly pathways are rather distinct and may therefore determine the fate of Nup145N. The C-terminal autoproteolytic domain (APD) of Nup145N and Nup116 can both recruit the cytoplasmic filament protein Nup82 *in vitro* (Fischer et al., 2015), but only engages with Nup116 *in vivo* (Kim et al., 2018). Our RIP-qPCR data reveal that Nup116 associates with the nascent chain of Nup82 (**Figure 2k**), suggesting that it determines Nup116 specificity for the cytoplasmic filaments. Taken together, our data highlight that not all motifs contained in linker Nups engage in co-translational interactions. We rather find that co-translational association events contribute to specifying paralogous assembly pathways (**Figure 2l**). Independent of our targeted analysis, none of the analyzed baits co-eluted their own mRNA in a co-translational manner (**Figure S2b**).

Inspired by the above findings, we wondered whether co-translational events may also specify assembly pathways for moonlighters that are members of multiple protein complexes. Two members of the Nup84-subcomplex, Sec13 and Seh1, are incomplete beta-propellers lacking one blade, that interact with the WD40 domain invasion motifs of Nup85 and Nup145C, respectively. While Seh1 and Sec13 both share moonlighting functions in the Seh1-associated (Sea) complex (**Figure S5b**) (Algret et al., 2014; Dokudovskaya et al., 2011), Sec13 further contributes to the architecture of the COPII coatomer complex (Fath et al., 2007; Whittle and Schwartz, 2010). We first investigated the domain invasion of Nup85 into the incomplete beta-propeller of Seh1 (**Figure 3a**) (Debler et al., 2008). RIP-qPCR analysis revealed that Seh1 enriched for the *nup85*-mRNA, but not the *sea4*-mRNA, in a translation-dependent manner (**Figure 3b**). To address if the co-translational association of full length Seh1 with the nascent chain of Nup85 is indeed mediated by the domain invasion motif at its N-terminus, we turned to SeRP. In line with our RIP-qPCR experiments, SeRP of Seh1-IPs identified a very strong enrichment of footprints within the *nup85*-ORF (**Figure 3c**). We analyzed the translatome-wide data but did not identify any similarly strong enrichment of footprints for any other gene (**Figure S5c-S5f**). This analysis suggests that the members of the Sea-complex which are known to interact with Seh1 (Algret et al., 2014; Dokudovskaya et al., 2011) but that remain to be structurally analyzed at high resolution, do not mimic assembly intermediates of Nup85 to an extent that warrants co-translational assembly. Surprisingly, the onset of the co-translational interaction of Seh1 with nascent Nup85 does not coincide with the emergence of the domain invasion motif (residue 44-101) from the exit channel of the ribosomes. It rather maps to alpha-helices and their connecting loops at positions 405-544. Although distant in sequence, these helices are located right beneath the domain invasion motif in the structure of the Seh1-Nup85 heterodimer (Stuwe et al., 2015b). Taken together, these findings illustrate that interaction motifs may co-translationally engage with some but not necessarily all of their moonlighting interaction partners and that the exact onset of co-translational interactions may adapt in a versatile way to protein folding. One may speculate that similar to the Sec13-Sec31 heterodimer (Čopič et al., 2012), interactions of Seh1 with the helical bundle may restrict the flexibility of the alpha-solenoid (**Figure 3d**) (see discussion).

**Figure 3:**
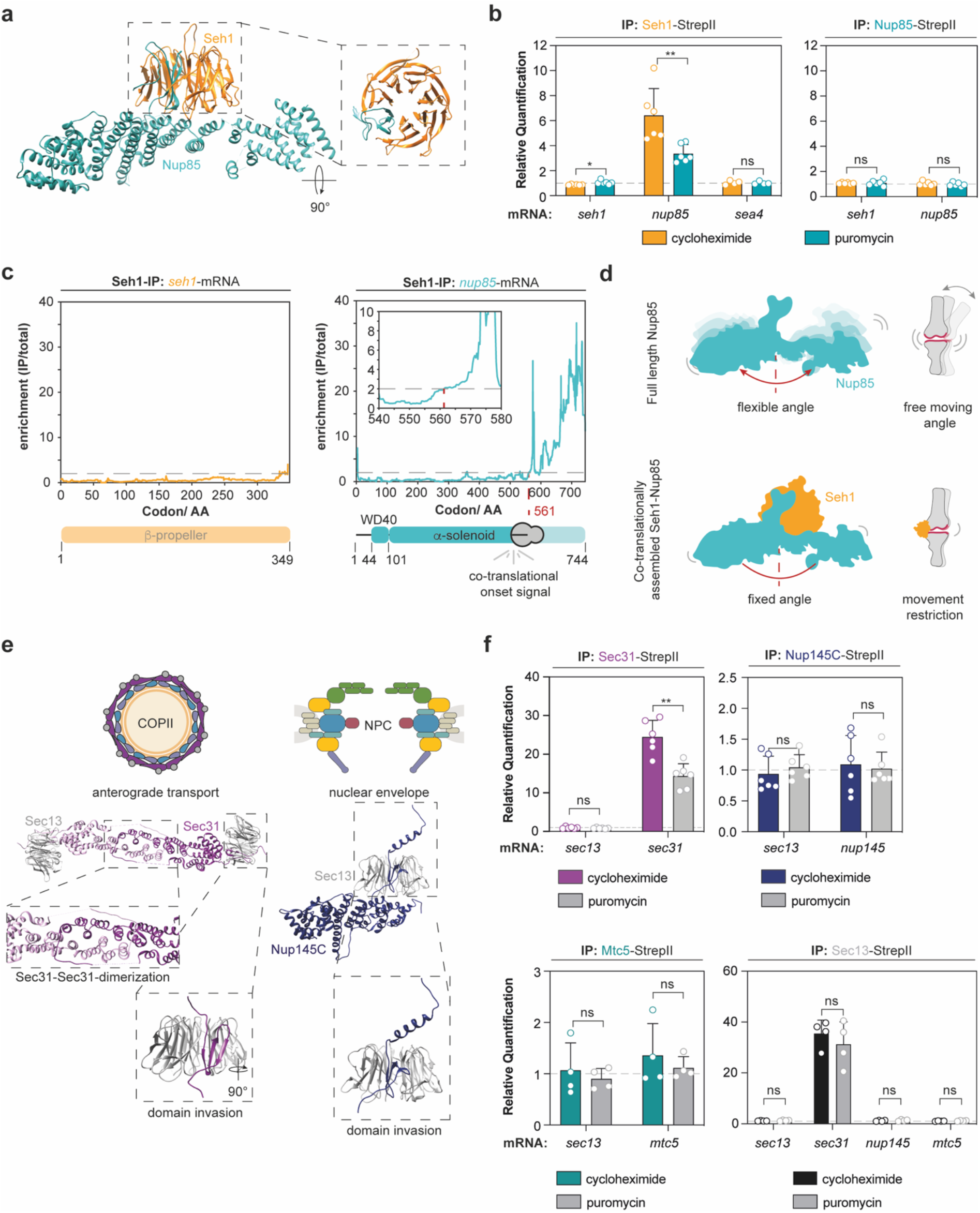
Beta-propellers can be complemented co-translationally *in vivo*. **a**, Crystal structure of Seh1 bound to Nup85 (PDB: 4XMM) (Stuwe et al., 2015b). **b**, RIP-qPCR reveals co-translational entanglement of Seh1 with Nup85. Bar graphs show mean ± SD of four to six biologically independent experiments. **c**, Selective ribosome profiling data derived from a Seh1-IP shows that helices at the trunk of Nup85 but not the domain invasion motif, are required for stable association with Seh1. Selective ribosome profiling was performed with four biologically independent replicates. **d**, Seh1 might restrict flexibility of the alpha-solenoid by binding to a structural joint represented by the helices it attaches to. **e**, Sec13 binds to the COPII cage by interacting with Sec31 (PDB: 2PM6) (Fath et al., 2007) and to the NPC by engaging with Nup145C (PDB: 3IKO) (Nagy et al., 2009) through beta-propeller complementation interactions. **f**, Co-translational interactions of Sec13 with any of the moonlighting interactors remained undetected, but Sec13 is bound to Sec31 when it co-translationally entangles with itself. Bar graphs depict mean ± SD of four to six biologically independent replicates. ns p>0.05,* p<0.05, ** p<0.01 (Two-sided, paired t-test). AA: amino acid; IP: immunoprecipitation; NPC: nuclear pore complex.

Sec13 was previously shown to engage with domain invasion motifs with at least three of the four known interactors (Algret et al., 2014; Dokudovskaya et al., 2011; Fath et al., 2007; Nagy et al., 2009; Whittle and Schwartz, 2010). Interestingly, Sec13 is present not only in the NPC and Sea-complex but also fulfills a dual role in COPII vesicles by interacting with Sec16 and Sec31 (**Figure 3e** and **S5b**). To identify the role of Sec13 for the assembly of the respective complexes, we purified Sec13, Sec31, Nup145C, and the Sea-complex protein Mtc5 and analyzed the co-eluted mRNAs by qPCR. We found that none of the purified components enriched for *sec13*-mRNA. However, RIP-qPCR revealed that Sec31 can co-translationally engage with its own mRNA. The IP against Sec13 strongly enriched for *sec31*-mRNA. However, puromycin treatment only weakly perturbed this interaction (**Figure 3f**). This finding may suggest that Sec13 binds to the *sec31*-mRNA directly, for example via its 3’UTR. Previously, it had been shown that protein binding to 3’UTRs can enforce interactions with nascent chains of orphan subunits which ultimately coordinate complex function and localization (Berkovits and Mayr, 2015; Lee and Mayr, 2019).

The Nsp1 complex is densely packed to the equatorial plane of the central channel of the inner ring. It consists of the Nsp1, Nup57, and Nup49 proteins. They contain N-terminal FG-rich intrinsically disordered domains that interact with nuclear transport receptors (Aramburu and Lemke, 2017). Each of the three members further contains a coiled-coil domain that heterotrimerizes thus forming the scaffold of the subcomplex (**Figure 4a**). The hetero-trimer embraces the IM-1 motif at the N-terminus of Nic96 (**Figure 2c**). This motif is thought to ensure the recruitment of the fully assembled CTN-complex to the inner ring of the NPC (Stuwe et al., 2015a). To elucidate if and how co-translational assembly events within the CTN-subcomplex contribute to the organization of the assembly pathway, we first studied the co-translational landscape by RIP-qPCR. Our results suggest that Nsp1 co-translationally binds to Nup57, but not Nup49 (**Figure 4b**). This is in line with previous *in vitro* analyses showing that Nsp1 cannot directly recruit Nup49 and hence Nup57 acts as an organizer subunit of the trimeric coiled-coil (Schlaich et al., 1997). To specify the onset of co-translational entanglement of Nsp1 with nascent Nup57 we used SeRP (**Figure 4c**). These experiments highlighted that the minimal requirement for co-translational interactions is the initial coiled-coil segment 1 (CCS1) that forms a rather long rod-shaped stretch. Neither RIP-qPCR nor selective ribosome profiling experiments detected a translation-dependent interaction of Nsp1 or Nup57 with the *nup49*-mRNA (**Figure 4b, 4c** and **S5g**) suggesting that Nup49 is added to the complex post-translationally. Furthermore, the C-terminal signal for Nup57 is rather surprising. Previously, most co-translational interactions were always detected at the N-terminus of the nascent chains (Shiber et al., 2018). Here, we show that the co-translational onset of not only Nup57 but also Nup85 can be positioned more towards to C-terminus and that those co-translational interactions might be warranted by a local decrease in translation efficiency (**Figure S6**).

**Figure 4:**
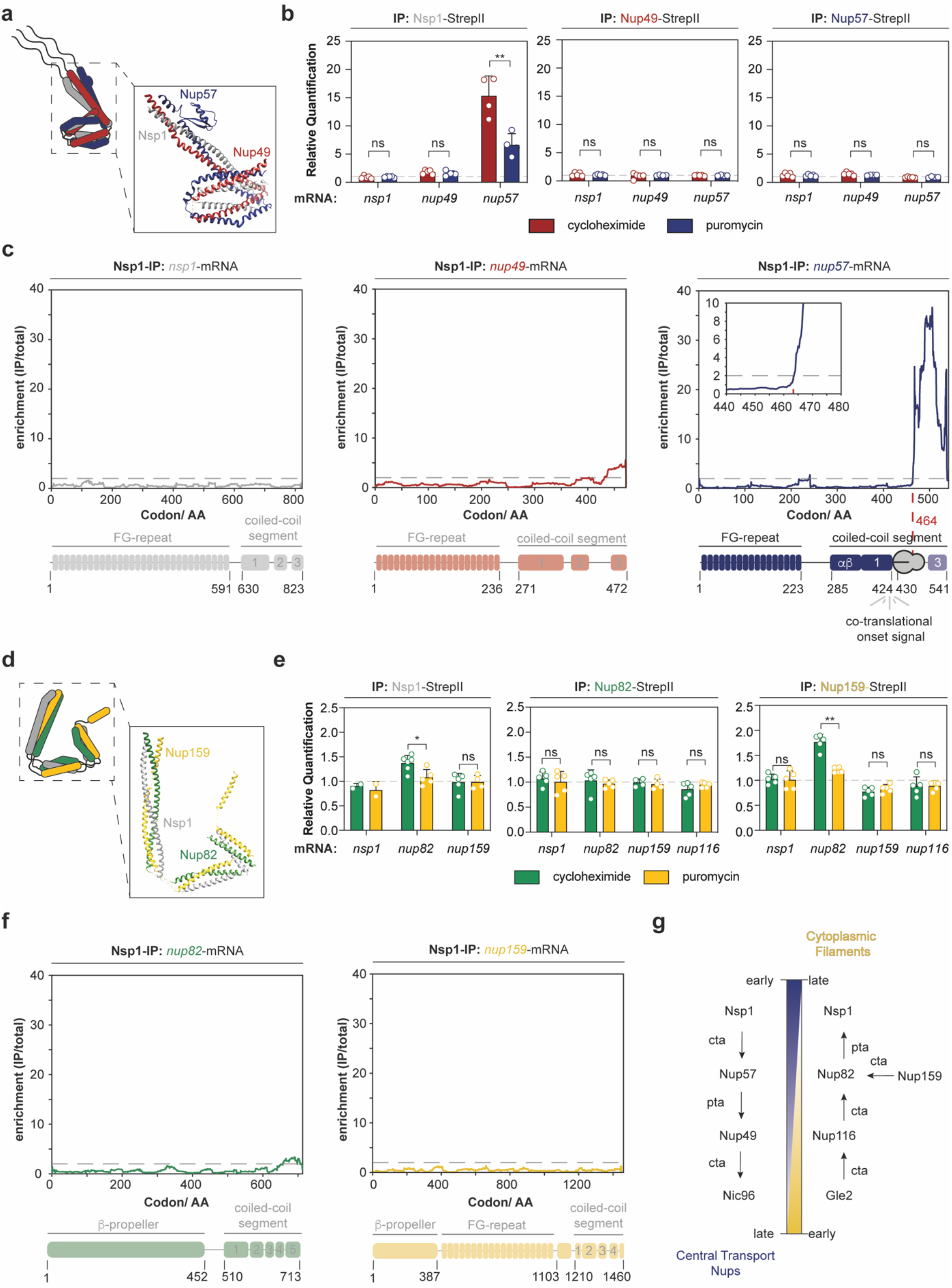
Nsp1 engages with the CTN and cytoplasmic filament subcomplexes in two opposing assembly pathways. **a**, Structure of *Chaetomium thermophilum* CTN shows the heterotrimeric coiled-coil which is tethered to Nic96 (PDB: 5CWS) (Stuwe et al., 2015a). **b**, RIP-qPCR of the CTN suggests co-translational interactions of Nsp1 with nascent Nup57 but not Nup49. Bar graphs show mean ± SD from four to six biologically independent replicates. **c**, Selective ribosome profiling from Nsp1-IPs identifies coiled-coil segment 1 (CCS1) as a fundamental asset for the co-translational association of Nsp1 with nascent Nup57. Selective ribosome profiling data was generated from four biologically independent replicates. **d**, Structural model of the coiled-coil in the Nup159-subcomplex (Kim et al., 2018) suggests an extended conformation. **e**, RIP-qPCR experiments targeting the Nup159-subcomplex (affinity purified Nsp1, Nup82 and Nup159). Nup159 co-translationally binds to nascent Nup82. RIP-qPCR experiments depicted as mean ± SD of two to six biologically independent replicates. **f**, SeRP with affinity purified Nsp1 does not detect co-translational association within the Nup159-subcomplex. Selective ribosome profiling was performed with four biologically independent replicates. **g**, Assembly scheme for the CTN- and Nup159-subcomplexes. Nsp1 co-translationally seeds the assembly in the CTN-subcomplex but post-translationally completes the assembly of the Nup159-subcomplex. ns p>0.05,* p<0.05, ** p<0.01 (Two-sided, paired t-test). IP: immunoprecipitation; AA: amino acid; cta: co-translational assembly; pta: post-translational assembly.

Intriguingly, the evolutionary related cytoplasmic filament subcomplex that functions as a platform for mRNA export (Weirich et al., 2004), is predicted to form a similar hetero-trimeric arrangement as the aforementioned CTN trimer (**Figure 4d**) (Gaik et al., 2015; Kim et al., 2018). It consists of Nup159, Nup82, and yet again Nsp1, whereby Nup57 and Nup82 compete for the same binding site within the Nsp1 coiled-coil region (Bailer et al., 2001). Due to competitive binding for Nsp1, we wondered if the cytoplasmic filaments are constructed by a similar assembly pathway. Surprisingly, RIP-qPCR and SeRP experiments targeting Nup82 and Nup159 showed no signal (**Figure 4e-f**, compare to **4b-c**). However, we found that Nup159 co-translationally engages with nascent Nup82 (**Figure 4e**), suggesting that in this case heterotrimer formation occurs in a reverse manner and Nsp1 is rather added post-translationally to the subcomplex. These findings suggest that the order of assembly of both subcomplexes helps to discriminate non-promiscuous interactions. Nsp1 binds to nascent Nup57 in a co-translational manner thus specifying the Nup57-Nsp1 dimer for interaction with full length Nup49 in the CTN-subcomplex. In the Nup159-subcomplex, however, the Gle2-Nup116-Nup159-Nup82 tetramer forms co-translationally first and only subsequently binds to Nsp1 (**Figure 4g**).

As we were surprised that the assembly pathways involving Nsp1 are so different, we queried which signal specifies the discrimination between Nup159- and CTN-subcomplex hetero-trimer formation. The sequences of the C-terminal coiled-coil segments of Nup57 and Nup82 align well with one another (**Figure 5a**). A notable exception, however, is a small alpha-beta domain upstream of the CCS1 that is expanded to a ferredoxin-like domain in vertebrates (Chug et al., 2015; Stuwe et al., 2015a). Deletion of this domain in *in vitro* reconstituted CTN-subcomplexes led to non-stoichiometric complexes (Stuwe et al., 2015a). We, therefore, hypothesized that the alpha-beta domain may set the stage for specific co-translational interactions of Nsp1 and Nup57 *in vivo*. To test whether this domain is required for co-translational assembly, we deleted the alpha-beta domain by scar-free cloning. Removal of the Nup57 unique alpha-beta domain did not impact thermo-sensitivity of the strain in respect to wildtype strain, showing that the deletion of the alpha-beta domain did not disrupt overall fitness (**Figure 5b**). Quantitative mass spectrometry analysis of Nsp1 pull downs from Nup57 wildtype and Nup57(Δαβ) strains revealed an enrichment of CTN components in the deletion strain, while the Nup159-subcomplex remained largely unaffected (**Figure 5c** and **S7**). Although alpha-beta-domain deletion perturbed the integrity of the CTN, it did not impair the co-translational assembly of Nsp1 - Nup57 or the CTN - Nic96 assemblome *in vivo* (**Figure 5d**), showing that the assembly defects occur post-translationally. Furthermore, the deletion did not cause a promiscuous co-translational signal for nascent Nup82 in RIP-qPCR analysis (**Figure 5d**). These findings indicate that the nascent CCS1 of Nup57 without the alpha-beta domain is still sufficient to be co-translationally discriminated from Nup82 and to bind to Nsp1.

**Figure 5:**
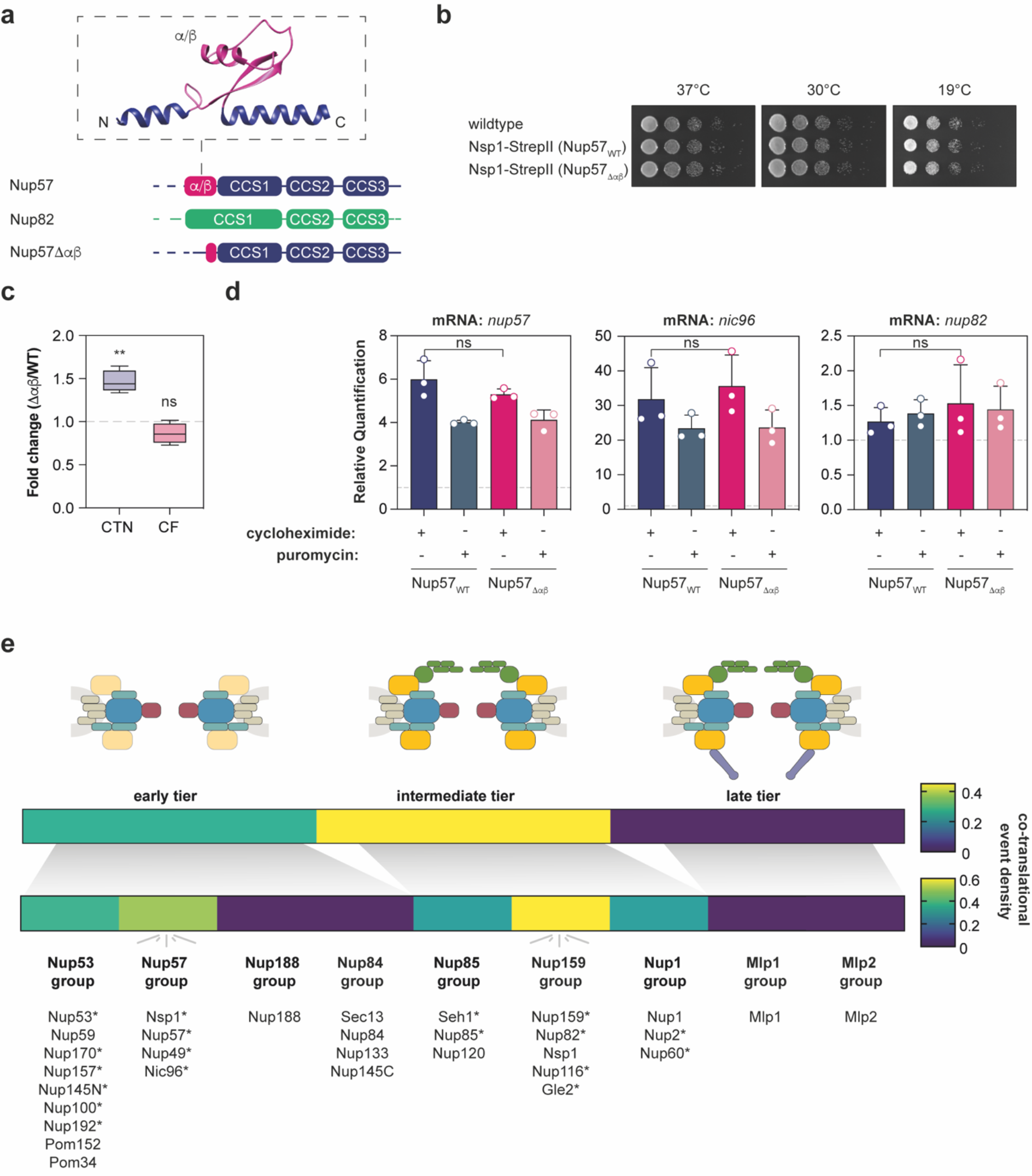
The alpha-beta domain of Nup57 does not specify for co-translational engagement with Nsp1. **a**, Structural scheme of the coiled-coil segments (CCS) of Nup82, Nup57 and Nup57Δαβ mutant. *Ct*Nup57 from PDB: 5CWS (Stuwe et al., 2015a). **b**, Growth phenotyping of Nup57 (wildtype) and Nup57Δαβ-mutant grown on YPD under permissive temperature. Deletion of the alpha-beta domain does not impair growth under permissive temperature. **c**, Visualization of quantitative mass spectrometry analysis of Nsp1-affinity purifications from wildtype or Nup57Δαβ strains. Fold-changes of CTN: central transport Nups (Nup57, Nup49) and CF: cytoplasmic filaments (Nup159, Nup82) reveal an enrichment for CTN but not CF components in the mutant strain, suggesting the formation of non-stoichiometric subcomplexes (dashed line: Nsp1 signal). Graph was generated from two biologically independent pull downs. **d**, RIP-qPCR analysis of Nsp1-IP experiments in wildtype and Nup57Δαβ-mutant. The mutant did not abolish co-translational interactions of Nsp1. Bar plots of RIP-qPCR data depict mean ± SD from three biologically independent replicates. e, Scheme visualizing the density of co-translational assembly events with respect to the order of the interphase assembly pathway as proposed by Onischenko et al. (2020). The heat map includes hits discovered in this study and previously published interactions (Lautier et al., 2021). Asterisk (*) marks co-translationally involved Nups. ns p>0.05,* p<0.05, ** p<0.01 (Two-sided, unpaired t-test).

## Discussion

Our hypothesis-driven approach has identified many co-translational assembly events that are relevant throughout all stages of NPC biogenesis with local hotspots at the subcomplexes utilizing Nsp1 (**Figure 5e**). Our qPCR experiments are in line with many, though not all interaction pairs reported by a recent screen (Lautier et al., 2021). Specifically, we identified translation-dependent interactions of Seh1 – *nup85*-mRNA, CTN – *nic96* mRNA, and Nsp1 - *nup57*-mRNA and Sec31 - *sec31*-mRNA, while we obtained negative data for Nup192 - *nup100*-mRNA. Differences may be attributed to the experimental design, in particular the use of translation-specific inhibitors such as cycloheximide and puromycin, the exact choice of biochemical conditions, and the design of scarlessly cloned yeast strains.

A very interesting discovery is that proteins moonlighting as parts of multiple complexes may co-translationally engage in one but not necessarily all alternative assembly pathways. This is exemplified by the selection of homologs by co-translational assembly pathways, such as Seh1 that does co-translationally interact with Nup85 but not with the Sea-complex and Nsp1 that does co-translationally interact with Nup57 but not with Nup82, at least under the conditions investigated here. Remarkably, the paralogous Nup159- and CTN-subcomplexes contain the most co-translational interactions. These data suggest that co-translational assembly may be used to seed ordered assembly pathways when multiple outcomes are possible and thus suppress promiscuous assembly.

We had initially expected that Sec13 may bind co-translationally to Nup145C, Sec31, and Mtc5, but found that Sec13 might rather associate with Sec31 post-translationally prior to the co-translational entanglement of Sec31 with itself. An interesting aspect is that the domain invasion motif of Nup85 is not sufficient for the co-translational interaction with Seh1, which requires the trunk of the supporting alpha-helical domain. Focusing on the importance of the helices, we speculate that Seh1 may rigidify the alpha-solenoid leading to a fully extended, slightly bent Nup85 (**Figure 3d**). This could be beneficial (i) to expose native interaction sites, for example, Nup120 (Kelley et al., 2015; Stuwe et al., 2015b) and/or (ii) to induce membrane curvature at the nuclear envelope. The latter benefit is supported by similar stiffening effects that were previously described for the Sec13-Sec31 heterodimer (Čopič et al., 2012). Here, an increase of rigidity was associated with the induction of membrane curvature of COPII vesicles and therefore might be reminiscent to the inside-out extrusion of the nuclear envelope in interphase assembly.

Seh1 also exemplifies that the onset of co-translational interactions may adapt in a versatile way to protein folding. In fact, it appears likely that the domain invasion motifs are subject to stronger selection pressure because its binding interfaces in the incomplete beta-propeller are the same for each interactor. Consequently, other structural features may be evolutionarily more accessible to organize a unique assembly pathway for the interactors of moonlighting proteins. The fact that minor changes in the open binding interfaces in the incomplete beta-propeller of Seh1 and Sec13 contribute to the selection of different binding partners was previously demonstrated by attempts to substitute Sec13 with Seh1 in a Sec31-Seh1 fusion construct within *sec13Δ*-*S. cerevisiae* strains. The fusion was shown to be lethal, highlighting the idea that the domain invasion blades are tailor-made for their native interactors (Čopič et al., 2012). One thus may speculate that the loops of i.e. Nup85 that engage with the surface of Seh1 may specify the interaction and provide additional affinity for its co-translational formation.

In our study, the CTN subcomplex nicely emphasizes the preservation of stoichiometry as another benefit of co-translational complex assembly. The CTN stoichiometry was highly controversial and had been addressed using different techniques (Chug et al., 2015; Melčák et al., 2007; Solmaz et al., 2011; Stuwe et al., 2015a; Ulrich et al., 2014). Our data point to a model where Nsp1 firstly binds the assembly domain of nascent Nup57, thus forming a 1 : 1 heterodimer. Next, full length Nup49 is bound in a post-translational manner into the 1 : 1 : 1 heterotrimer, which subsequently binds to the IM-1 assembly domain of Nic96, yet again co-translationally, resulting in a 1 : 1 : 1 : 1 hetero-tetramer. Although the deletion of alpha-beta domain of Nup57 showed stoichiometric defects for CTN components (Nup49 and Nup57), the effect size at this stage was not as severe as for *in vitro* reconstituted complex (Stuwe et al., 2015a) and stoichiometry of Nic96 remained unaffected, showing that the two co-translational events may warrant that only stoichiometric subcomplex can be implemented into NPCs. This experimentally determined that the linear assembly outline is strikingly in line with our theoretical considerations (**Figure 1**), and shows that not necessarily all, but at least several of the individual steps of a given assembly pathway may occur co-translationally. Taken together, our findings provide new insights into NPC biogenesis. It has been previously shown that most of the Nup-subcomplex encoding mRNAs, including those of the CTN complex (Nup62 in *Drosophila melanogaster*), are translated without any detectable co-localization on the mRNA level (Hampoelz et al., 2019b). The advantage of translation under relatively dilute conditions in the cytosol could be that stoichiometric subcomplexes are formed but higher-order interactions, such as subcomplex oligomerization, are prevented due to the low protein concentrations. The benefit of co-translational assembly could be to increase the dwell-time of assembly intermediates to nevertheless facilitate efficient assembly. Hereby, at least four early modules are synthesized separately, the Nup84-, Nsp1-, Nup159- and inner ring complexes, the latter consisting of Nup53-Nup170 and Nup157-Nup145 pairs. This is in line with recent mass spectrometric studies that have revealed the order of subunit engagement of NPC biogenesis (Onischenko et al., 2020).

Next, subcomplexes are recruited to sites of NPC biogenesis in proximity to membranes where their local concentration is increased and higher-order interactions across subcomplexes, such as ring formation, become kinetically favored. Here, Nup53 and Nup170 that were previously assembled co-translationally seed the recruitment of further components such as Nup188 and Nup192 (Onischenko et al., 2020) presumably already associated with Nup100 (Lautier et al., 2021). Subsequently, additional subcomplexes are recruited, various of which were pre-assembled stoichiometrically in a co-translational manner. The concept of co-translational assembly thus elegantly complements present scientific models of NPC biogenesis and explains how promiscuous interactions are avoided in such a very complex macromolecular assembly.

## Authors Contribution

M.S. conceived the project, designed experiments, performed experiments, analyzed data, and wrote the manuscript. A.B. performed experiments. F.P. designed experiments. J.J.M.L. and N.T.D.d.A performed experiments, analyzed data, and wrote the manuscript. C.M.F. performed experiments. E.K. performed experiments. J.B. designed experiments, analyzed data, wrote the manuscript. J.D.L., E.M.S., and K.R.P. supervised the project. G.H. performed modeling and wrote the manuscript. V.B. designed experiments, supervised the project. M.B. conceived the project, designed experiments, analyzed data, supervised the project, and wrote the manuscript.

## Acknowledgment

We thank Patrick Hoffmann, Natalie Romanov, and Florian Wilfling for the fruitful discussion on the manuscript. Additionally, the authors like to thank Georg Stoecklin, Lars Steinmetz, and Britta Brügger for the critical assessment of the project. M.B. acknowledges funding by the Max Planck Society and the European Research Council (724349-ComplexAssembly).

## Materials and Methods

### 1 Yeast Strains and Growth Media

StrepII-tagged yeast strains were obtained by scar-free homologous recombination using the MX4 blaster cassette (Carvalho et al., 2013). Briefly, the MX4 blaster cassette was amplified with gene-specific overhangs for homologous recombination. PCR products were transformed and positive clones were selected on YPD-high phosphate plates supplemented with 300 μg/mL hygromycin B (ForMedium) and 3 g/l potassium dihydrophosphate (monobasic). To remove the MX4 blaster cassette, MX4 positive clones were grown in low phosphate YPD to induce endonuclease expression (Carvalho et al., 2013) and were transformed with a codon-optimized twin-StrepII-tag (5’ TCTGCTTCTGCTTGGTCACATCCACAATTTGAAAAAGGTGGTGGTTCTGGTGGCGGTTCAGGTGGTTCATCTGCTTGGAGTCATCCTCAATTCGAAAAG 3’) (Schmidt et al., 2013) with respective gene-specific overhangs. Once transformed, clones were screened on YP-galactose plates. The successful transformation was validated by PCR.

For Sec13, the transformation of the MX4 blaster cassette remained unsuccessful unless an additional copy of Sec13 was inserted into a pRS423 overexpression plasmid. To obtain MX4 positive clones, the aforementioned strategy was used. To select the clones, yeast was plated onto His-drop out plates (ForMedium) containing 1 g/L mono-sodium glutamate, 1.9 g/L YNB without amino acids and ammonium sulfate (ForMedium), 3 g/L potassium dihydrophosphate (monobasic), and 300 μg/mL hygromycin B. To insert the StrepII-tag, yeast strains were propagated in the aforementioned media to trigger the loss of the overexpression plasmid. The insertion of the C-terminal StrepII-tag was validated by PCR. Additionally, StrepII-positive strains were plated on His-Drop out plates to ensure *HIS*-marker removal.

For growth phenotyping, yeast strains were incubated overnight in YPD. On the next day, OD(600) was determined and set to OD(600) of 1 using YPD. Serial dilution in a ratio of 1:10 was prepared in YPD and spotted onto YPD plates. Strains were grown at indicated temperatures.

### 2 RIP-qPCR experiments

The protocol for the RIP-qPCR experiments is an adaptation of the previously published methods (Kamenova et al., 2019; Shiber et al., 2018). For RIP-qPCRs, overnight cultures were grown in YPD. These cultures were used to set 400 mL of YPD to an OD(600) of 0.035. The expression cultures were grown at 30°C, 160 rpm, and cultured to an OD(600) of 0.5-0.6. Then, cultures were harvested by rapid filtration onto nitrocellulose membrane (0.45 μm; Bio-Rad) and cells were flash-frozen in liquid nitrogen.

Once frozen, cells were supplemented with 1.4 mL of frozen high salt lysis buffer (20 mM Hepes-KOH, pH 7.5, 500 mM KCl, 20 mM MgCl_2_, 1 mM PMSF, 0.01 % IGEPAL and cOMPLETE EDTA-free protease inhibitor (Roche), 0.1 mg/mL CHX (Sigma-Aldrich) or 0.01 mg/mL puromycin (Sigma-Aldrich)). Cells were disrupted under cryogenic conditions using the CryoMill (Retsch) at 30 Hz for 2 min.

The lysate was thawed and transferred into 1.5 mL tubes. The crude lysate was cleared at 15,000 *g* at 4°C for 3 min. Afterward, the cleared supernatant was loaded onto equilibrated Streptactin resin (IBA) supplemented with 60 μL of BioLock (IBA) to prevent unspecific binding and 0.1 U/μL Ribolock (Invitrogen) to inhibit RNA decay. The lysate was incubated on the beads by end-to-end mixing at 4°C for 1 hr. Then, beads were subjected to subsequent wash steps. Briefly, beads were centrifuged at 500 *g* and 4°C for 5 min. Supernatant was removed and beads were washed 3-times with 1 mL of wash buffer A (20 mM Hepes-KOH, pH 7.5, 140 mM KCl, 20 mM MgCl_2_, 0.01 % IGEPAL and cOMPLETE EDTA-free protease inhibitor, 0.1 mg/mL CHX or 0.01 mg/mL puromycin) for 1 min by end-to-end mixing, followed by two washes (1 min, 4 min) with wash buffer B (20 mM Hepes-KOH, pH 7.5, 140 mM KCl, 20 mM MgCl_2_, 0.05 % IGEPAL and cOMPLETE EDTA-free protease inhibitor, 0.1 mg/mL CHX or 0.01 mg/mL puromycin). After the washes, the beads were resuspended in 500 μL of 10 mM Tris-HCl, pH 8.0.

RNA was extracted by adding 40 μL of 20 % SDS and the addition of 750 μL of pre-warmed phenol-chloroform-isoamyl alcohol (PCI, 65°C, Invitrogen). This mixture was then incubated at 65°C, 1400 rpm for 5 min followed by snap cooling on ice for 10 min. Next, extractions were centrifuged at 15,000 *g* for 10 min and the aqueous phase was again subjected with 750 μL of PCI. This time, the extraction was performed at room temperature and occasional vertexing for 5 min. The centrifugation was repeated. Finally, residual PCI was removed by a diethyl ether wash, and the remaining organic solvent was evaporated in a Speedvac (Eppendorf). RNA was precipitated by adding 3 M NaOAc, pH 5.5 to reach a final concentration of 0.3 M, 2.5 μL Glycoblue (Invitrogen), and equivalent amounts of isopropanol. Precipitates were placed into the −80°C freezer overnight. Samples were centrifuged at 15,000 *g* and 4 °C for 90 min. The resulting pellet was washed in 70 % EtOH, dried in a Speedvac (Eppendorf), and resuspended in 20 μL of 10 mM Tris-HCl, pH 8.0. Typically, precipitations yielded 150-250 ng/μL of RNA.

For reverse transcription, 500 ng of RNA were applied and cDNA was synthesized according to the manufacturer’s instructions of the VILO Reverse Transcription kit (Invitrogen) including the optional ezDNase step.

Real-time qPCR was conducted using the TaqMan Fast and Advanced Master Mix (Applied Biosystems) according to the manufacturer’s manuscript. FAM-labelled qPCR probes were purchased from Applied Biosystems as specified in **Table 1**. The qPCR was performed using the QuantStudio 5 cycler (Applied Biosystems, 50°C: 2 min, 95°C: 2min; 40 cycles: 95°C: 0:01 min, 60°C: 0:20 min). Images were taken every cycle within the annealing/extension step. All qPCR assays were performed in technical triplicates and each experiment was analyzed using the QuantStudio analysis software (v1.5.1).

**Table 1:** TaqMan probes used for RIP-qPCR experiments.

| Gene | TaqMan Probe Name | Fluorophore | Supplier |
| --- | --- | --- | --- |
| *act1* | Sc04120488_s1 | FAM-MGB | Applied Biosystems |
| *seh1* | Sc04122707_s1 | FAM-MGB | Applied Biosystems |
| *nup85* | Sc04138225_s1 | FAM-MGB | Applied Biosystems |
| *nsp1* | Sc04135209_s1 | FAM-MGB | Applied Biosystems |
| *nup49* | Sc04123659_s1 | FAM-MGB | Applied Biosystems |
| *nup57* | Sc04126036_s1 | FAM-MGB | Applied Biosystems |
| *nic96* | Sc04120807_s1 | FAM-MGB | Applied Biosystems |
| *nup145* | Sc04122572_s1 | FAM-MGB | Applied Biosystems |
| *sec31* | Sc04108310_s1 | FAM-MGB | Applied Biosystems |
| *sec13* | Sc04147734_s1 | FAM-MGB | Applied Biosystems |
| *nup82* | Sc04135552_s1 | FAM-MGB | Applied Biosystems |
| *nup159* | Sc04133316_s1 | FAM-MGB | Applied Biosystems |
| *nup116* | Sc04153403_s1 | FAM-MGB | Applied Biosystems |
| *nup100* | Sc04140578_s1 | FAM-MGB | Applied Biosystems |
| *nup53* | Sc04154666_s1 | FAM-MGB | Applied Biosystems |
| *nup170* | Sc04099596_s1 | FAM-MGB | Applied Biosystems |
| *nup157* | Sc04118666_s1 | FAM-MGB | Applied Biosystems |
| *nup192* | Sc04135187_s1 | FAM-MGB | Applied Biosystems |
| *gle2* | Sc04118682_s1 | FAM-MGB | Applied Biosystems |
| *sea4* | Sc04100019_s1 | FAM-MGB | Applied Biosystems |
| *mtc5* | Sc04110895_s1 | FAM-MGB | Applied Biosystems |
| *fas1* | Sc04141945_s1 | FAM-MGB | Applied Biosystems |
| *fas2* | Sc04172723_s1 | FAM-MGB | Applied Biosystems |

### 3 Selective Ribosome Profiling

The selective ribosome profiling experiments were conducted according to previously published protocols by the Bukau lab (Galmozzi et al., 2019; Shiber et al., 2018). Each biologically independent replicate was obtained from 800 mL of yeast cultures grown in YPD to an OD(600) of 0.5-0.6 in analogy to the RIP-qPCR experiment. Harvest was performed as previously described for the RIP-qPCR experiments. Subsequently, cells were lysed in 3 mL of ribosome profiling buffer (20 mM Hepes-KOH pH 7.5, 140 mM KCl, 10 mM MgCl_2_, 1 mM PMSF, 0.01 % IGEPAL, 0.1 mg/mL CHX, 1 tablet of cOMPLETE protease inhibitor per 50 mL) using the cryo-mill (30 Hz, 2 min).

Then, the lysate was thawed and centrifuged at 15,000 *g* at 4°C for 3 min. The absorbance at 260 nm of a 1:100 dilution was measured to determine the amount of RNase I (Ambion). Here, 20 U of RNase I per A260 was applied to convert polysomes into monosomes. RNase I was incubated by end-to-end mixing for 20 min at 4°C. The reaction was quenched by adding 200 U Superase·In (Invitrogen). Then, ribosomes were pelleted using a 25 % sucrose cushion (20 mM Hepes-KOH pH 7.5, 140 mM KCl, 10 mM MgCl_2_, 25 % w/v sucrose, 0.01 % IGEPAL, 0.1 mg/mL CHX, 1 tablet of cOMPLETE protease inhibitor per 50 mL) at 150,000 *g* for 2.5 hr. The ribosomal pellets were resuspended in 1 mL of wash A containing 10 U DNase I (RNase-free, Thermo Scientific) and 30 μL BioLock (IBA). 100 μg of RNA was taken for the total translatome library. The remainders were applied to 250 μL of pre-equilibrated Streptactin sepharose (IBA). The pull down and RNA precipitation was performed as stated in the RIP-qPCR experiments.

Precipitated RNA was resuspended in 20 μL of TE buffer and supplemented with equal amounts of 2 x RNA loading dye (Thermo Scientific). In parallel, the RNA marker was prepared by mixing the low range RiboRuler (Thermo Scientific) with 200 nM synthetic 5’FAM labeled 34-mer, 30-mer, 28-mer and 26-mer. Sequences of the custom made RNAs are listed in **Table 2**.

**Table 2:**
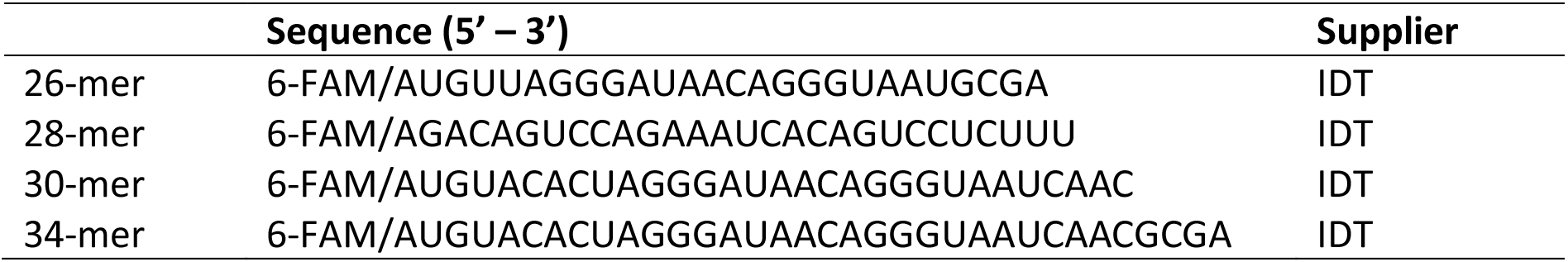
Synthetic RNAs (adapted from (McGlincy and Ingolia, 2017; Welz, 2003)) were used as markers for size selection of ribosome footprints. For detection reasons, RNA was fluorophore-labeled with 6-FAM on the 5’ end.

Both were denatured at 80°C for 2 min and put back on ice. 15 % denaturing PAGE (Carl Roth) was prepared, pre-warmed for 1 hr at 16 W and then loaded. The gels were run at 16 W for 3.5-4 hr until the bromophenol blue emerged. Afterward, the gels were stained using SybrGold (Invitrogen) and imaged using the Amersham Typhoon (GE Healthcare). The area between 26-34 nt were excised and crushed. The RNA was eluted in 500 μL of Tris-HCl pH 8.0 at 70°C for 10 min while shaking at 14,000 rpm. Elutant was separated from gel pieces by putting them through a Spin-X cellulose acetate column (Corning). The RNA was precipitated by supplying 50 μL 3 M NaOAc pH 5.5, 2.5 μL Glycoblue co-precipitation agent (Invitrogen), and 500 μL isopropanol.

The purified RNA was initially dephosphorylated in 1 x FastAP buffer containing 2 U FastAP (Thermo Scientific) and 20 U RiboLock (Invitrogen). The reaction was incubated for 15 min, at 37°C, and 600 rpm and immediately heat-inactivated at 75 °C for 5 min. 5’ ends were phosphorylated by 20 U polynucleotide kinase (PNK; NEB) by adding 1 mM ATP (Thermo Scientific), 1 x PNK buffer (NEB), and 20 U RiboLock. The reaction was incubated for 30 min at 37 °C.

Afterward, RNA integrity and concentration were checked using the RNA Pico 6000 Assay Kit of the Bioanalyzer 2100 system (Agilent Technologies). Small RNA libraries were prepared from 1 ng of RNA using the NEXTflex Small RNA-seq Kit v3 (Perkin Elmer). The size distribution of the libraries was assessed on a Bioanalyzer with a DNA High Sensitivity kit (Agilent Technologies), and concentration was measured with the Qubit DNA High Sensitivity kit in the Qubit 2.0 Fluorometer (Life Technologies). Subsequently, libraries that passed the QC step were pooled in equimolar amounts and the final pool was purified with SPRI select beads with a 1.3x ratio (Beckman Coulter). The final pool was loaded on the Illumina sequencer NextSeq500 High output and sequenced uni-directionally, generating ^~^500 million reads, each 85 bases long.

### 4 Polysome Profiling

5% and 45 % sucrose (w/v) was dissolved in 20 mM Hepes-KOH pH 7.5, 140 mM KCl, 10 mM MgCl_2_, 0.01 % IGEPAL, 0.1 mg/mL CHX and 1 tablet of cOMPLETE protease inhibitor per 50 mL. Gradients were mixed in thin-wall polypropylene tubes (Beckman, 331372) using a gradient mixer (BioComp) and equilibrated overnight at 4°C. RNA concentration of the cleared lysate was measured by nanodrop and 500 μg of this RNA was loaded onto the gradient and run for 2.5 hr at 220,000 *g* and 4°C in an SW41-rotor (Beckman). Gradients were then run at 850 μL/min in a density gradient fractionation system (Teledyne Isco), chased by 60 % sucrose in water. RNA absorbance at 254 nm was continuously measured using a UA-6 detector with the sensitivity setting 2.

### 5 Analysis of Nsp1-subcomplexes

1 L of yeast culture was set to an OD(600) 0.05 and grown to OD(600) 1.2-1.4 at 30°C and 130 rpm in baffle flasks. Cells were harvested by centrifugation, washed in ice-cold PBS and resuspended in Hepes-NB (20 mM Hepes pH 7.5, 150 mM NaCl, 50 mM K(OAc), 2 mM Mg(OAc)_2_, 1 mM DTT, 5 % glycerol, 0.01 % (v/v) IGEPAL, 1 mM benzamidine, 1 tablet/50 mL of cOmplete EDTA-free protease inhibitor and 1 mL/50 mL BioLock (IBA)) in adaptation to Fischer et al. (2015). Resuspension was frozen drop-wise in liquid nitrogen and lysed using the cryo-mill (30 Hz, 2 min).

The cell lysate was thawed and cell debris was pelleted by ultracentrifugation (35,000 *g*, 20 min, 4°C). The supernatant was applied to 500 μL bed volume of Streptactin sepharose resin and incubated for 1 hr on a rolling mixer at 4°C. The resin was washed with 4 x 5 mL of Hepes-NB. Protein was eluted in Hepes-NB supplemented with 20 mM D-desthiobiotin (IBA) in four elution steps (3 x 350 μL and 1 x 500 μL) each time incubating the resin 5 min with the elution buffer.

To avoid unnecessary dilution of the elution fractions, the first fraction was omitted. After elution, protein concentration was determined by measuring A280. 500 μL of elution was immediately supplemented with 20 mM TCEP and incubated at 30 min at 37°C. Next, the pull downs were subsequently alkylated using 20 mM iodoacetamide (IAA) incubated in the dark for 20 min at room temperature and further processed by adding 12 % aqueous phosphoric acid to obtain a final concentration of 1.2 % of phosphoric acid.

Pre-processed pull downs were mixed with S-trap binding buffer, transferred to S-trap ProtiFi plates (ProtiFi), and treated according to the manufacture’s protocol. Finally, the protein was converted into peptides using a 1:100 Trypsin : protein ratio by supplementing the corresponding amount of Trypsin in 125 μL of digestion buffer that was added to each condition. Trypsin digest was carried out overnight at 4°C.

Before elution, 80 μL of digestion buffer was added to each well of the S-trap digestion plate and eluted in an OASIS elution plate (Waters). Next, 80 μL of 0.2 % of aqueous formic acid was added per well and elution was repeated. Finally, 80 μL of aqueous acetonitrile (ACN) containing 0.2 % formic acid was applied and peptides were recovered. The eluted peptides were transferred and solvents were evaporated in a speed vac (Eppendorf). Dried peptides were resolved in 80 μL of HPLC water. 20 μL of these peptides were then subjected for peptide concentration assays (Thermo Scientifc). The remaining peptides were cleaned up using the OASIS desalting plates.

Dried peptides were reconstituted in 5% acetonitrile (ACN) with 0.1% formic acid (FA). Peptides were loaded onto a C_18_-CoAnn trapping column (particle size 3 μm, L = 20 mm) and separated on a C_18_-CoAnn analytical column (particle size = 2 μm, ID = 75 μm, L = 50 cm, CoAnn Technologies, LLC, Richland, USA) using a nano-HPLC (Dionex U3000 RSLCnano) at a temperature of 55°C.

Trapping was carried out for 6 min with a flow rate of 6 μL/min using a loading buffer (100 % H_2_O with 0.05 % trifluoroacetic acid). Peptides were separated by a gradient of water (buffer A: 100 % H_2_O and 0.1 % FA) and acetonitrile (buffer B: 80 % ACN, 20 % H_2_O, and 0.1 % FA) with a constant flow rate of 250 nL/min. The gradient went from 4 % to 48 % buffer B in 90 min. All solvents were LC-MS grade and purchased from Riedel-de Häen/Honeywell (Seelze, Germany).

Eluting peptides were analyzed in data-dependent acquisition mode on a Fusion Lumos mass spectrometer (ThermoFisher Scientific) coupled to the nano-HPLC by a Nano Flex ESI source. MS1 survey scans were acquired over a scan range of 350 to 1400 mass-to-charge ratio (m/z) in the Orbitrap detector (resolution = 120k, automatic gain control (AGC) = 2e5, and maximum injection time: 50 ms). Sequence information was acquired by a “ddMS2 OT HCD” MS2 method with a fixed cycle time of 2 s for MS/MS scans. MS2 scans were generated from the most abundant precursors with a minimum intensity of 3e4 and charge states from two to five. Selected precursors were isolated in the quadrupole using a 1.4 Da window and fragmented using higher-energy C-trap dissociation (HCD) at 30 % normalized collision energy. For Orbitrap MS2, an AGC of 1e4 and a maximum injection time of 54 ms were used (resolution = 30k). Dynamic exclusion was set to 30 s with a mass tolerance of 10 parts per million (ppm). Each sample was measured in duplicate LC-MS/MS runs.

MS raw data were processed using the MaxQuant software (v1.6.6.0) with customized parameters for the Andromeda search engine. Spectra were matched to a *Saccharomyces cerevisiae* database downloaded from UniProtKB (April 2021), a contaminant and decoy database, with a minimum Tryptic peptide length of seven amino acids and a maximum of two missed cleavage sites. Precursor mass tolerance was set to 4.5 ppm and fragment ion tolerance to 20 ppm, with a static modification (carboxyamidomethylation) for cysteine residues. Acetylation on the protein N-terminus and oxidation of methionine residues were included as variable modifications. A false discovery rate (FDR) below 1 % was applied at protein, peptide, and modification levels. The “match between runs” option was enabled and only proteins identified by at least one unique peptide were considered for further analysis. All proteomics data (including acquisition and data analysis parameters) associated with this manuscript have been deposited at the ProteomeXchange Consortium (http://proteomecentral.proteomexchange.org) via the PRIDE partner repository (Perez-Riverol et al., 2019). For revision, anonymous reviewer account credentials are available upon request.

Abundance changes were calculated from median-centered peptide intensities. The mean of median-centered peptide intensities was normalized to the intensity of Nsp1 within the same biological replicate and a fold change was calculated. For **Figure S7**, the mean of peptide intensities were calculated and normalized to Nsp1.

### 6 Protein Analysis

4 μL of the crude lysate (representing 0.4 %) in 1 x NuPAGE loading dye (Invitrogen) and 10 μL of boiled beads (representing 4 %) in 1 x NuPAGE loading dye were loaded onto NuPAGE Bis-Tris gels (MW<100 kDa) (Invitrogen) or Tris-Glycine gels (Bio-Rad; MW>100 kDa) and run at 160 V for 50 min. Protein was transferred onto 0.45 μm TransBlot Turbo nitrocellulose (Bio-Rad) using the High MW setting of the TurboBlot system (Bio-Rad) according to the manufacturer’s procedure. Membranes were blocked in 5 % milk in TBS-T (0.02 % Tween-20) for 1 hr at room temperature under gentle shaking. Primary antibody (1:5000) was added and incubated overnight at 4°C while constant shaking. Next, membranes were washed in TBS-T and a secondary antibody (1:10,000) was applied. The membrane was stained for 1 hr at room temperature. Before visualization by ECL developing solution (Bio-Rad), membranes were washed again. Membranes were imaged using the Chemidoc (Bio-Rad). Antibodies used in this study are listed in **Table 3**.

**Table 3:** List of antibodies used in this study.

|  | Supplier | Identifier |
| --- | --- | --- |
| Recombinant Anti-Strep tag II antibody | abcam | EPR12666; ab180957 |
| Anti-rabbit IgG, HRP-linked Antibody | Cell Signaling Technology | x |

Fas1 and Fas2 IPs were stained using Instant Blue (abcam) according to the manufacturer’s protocol. For Nsp1 pull downs, NuPAGE Bis-Tris gels were stained using the SilverQuest Silver Staining Kit (Invitrogen).

### 7 Translation Efficiency Profiles

Normalized translation efficiency profiles for *S. cerevisiae* were calculated using a previously published pipeline by Frydman lab (https://web.stanford.edu/group/frydman/codons/codons.html) (Pechmann and Frydman, 2013).

### 8 Data Processing

Sequencing reads were processed according to the guidelines published in Galmozzi et al., 2013 (Galmozzi et al., 2019). In brief, reads were cleaned and trimmed using cutadapt (v2.3) (Martin, 2011). Sequences mapping to *Saccharomyces cerevisiae* noncoding RNA (R64-1-1.ncrna) were discarded and the remaining reads were mapped to *Saccharomyces cerevisiae* (R64-1-1) genome using tophat2 (v2.0.10). From the script suite (https://doi.org/10.5281/zenodo.2602493), only supplementary script A was applied on each sample that were then used to assign ribosome positions.

The number of reads per genomic position was extracted using script A and was used as input for subsequent analysis using in-house MATLAB scripts (v9.7.0.1296695 (R2019b) Update 4). The scripts combined data from different replicates and used gliding averages to evaluate the enrichment within a given sequence window.

### 9 Statistics and data analysis

Data in figures was illustrated as mean with corresponding standard deviation (SD) using GraphPad Prism (v9.0.0). Dashed lines in qPCR graphs represent wildtype background levels (no bait) determined for cycloheximide and puromycin, respectively. Significance levels of qPCRs for one mRNA obtained from the same bait under cycloheximide and puromycin treatments was determined by applying a two-sided Student’s t-test for paired samples by assuming a normal distribution of the data unless otherwise stated (ns: p > 0.05; *p< 0.05; **p< 0.01; ***p <0.001). Comparison of Nup57(wildtype) and Nup57(Δαβ)-strains was conducted using a two-sided, unpaired t-test (ns: p > 0.05; *p< 0.05; **p< 0.01; ***p <0.001). For mass spectrometry, Nup49 and Nup57 and Nup159 and Nup82 were grouped into CTN and cytoplasmic filaments, respectively. Significance levels were calculated in respect to Nsp1 by a two-sided, unpaired Student’s t-test (ns: p > 0.05; *p< 0.05; **p< 0.01; ***p <0.001). Co-translational event density (**Figure 5e**) was calculated by dividing the number of co-translational events by the number of proteins per group. Secondary binding and collective binding as previously observed for the translation dependent interactions of Gle2 - *nup82* (Lautier et al., 2021) and CTN - *nic96* was considered as one event. Structures were analyzed using UCSF Chimera (v1.15).

### 10 Strains

Yeast strains used in this study are listed in **Table 4**.

**Table 4:** Yeast strains and corresponding genotypes used in this study.

| **Strain Name** | **Genotype** | **Source** |
| --- | --- | --- |
| BY4741 (wt) | *MATa his3Δ1 leu2Δ0 met15Δ0, ura3Δ0* | provided by Patil lab |
| Nic96-StrepII | (BY4741)*nic96-strepII* | this study |
| Nsp1-StrepII | (BY4741)*nsp1-strepII* | this study |
| Nup49-StrepII | (BY4741)*nup49-strepII* | this study |
| Nup57-StrepII | (BY4741)*nup57-strepII* | this study |
| Nup53-StrepII | (BY4741)*nup53-strepII* | this study |
| Nup192-StrepII | (BY4741)*nup192-strepII* | this study |
| Nup157-StrepII | (BY4741)*nup157-strepII* | this study |
| Nup170-StrepII | (BY4741)*nup170-strepII* | this study |
| Nup100-StrepII | (BY4741)*nup100-strepII* | this study |
| Nup116-StrepII | (BY4741)*nup116-strepII* | this study |
| Gle2-StrepII | (BY4741)*gle2-strepII* | this study |
| Seh1-StrepII | (BY4741)*seh1-strepII* | this study |
| Nup85-StrepII | (BY4741)*nup85strepII* | this study |
| Sec13-StrepII | (BY4741)*sec13-strepII* | this study |
| Sec31-StrepII | (BY4741)*sec31-strepII* | this study |
| Nup145C-StrepII | (BY4741)*nup145-strepII* | this study |
| Mtc5-StrepII | (BY4741)*mtc5-strepII* | this study |
| Nup82-StrepII | (BY4741)*nup82-strepII* | this study |
| Nup159-StrepII | (BY4741)*nup159-strepII* | this study |
| Fas1-StrepII | (BY4741)*fas1-strepII* | this study |
| Fas2-StrepII | (BY4741)*fas2-strepII* | this study |
| Nsp1-Strep (Nup57$\Delta\alpha\beta$) | (BY4741)*nsp1-strepII, nup57(Δ925-1062)* | this study |

