## Supplementary Figures S1-S7 for "Co-translational assembly counteracts promiscuous interactions"

**The supplementary file includes:**

Figs. S1 to S7

### Supplementary Figures

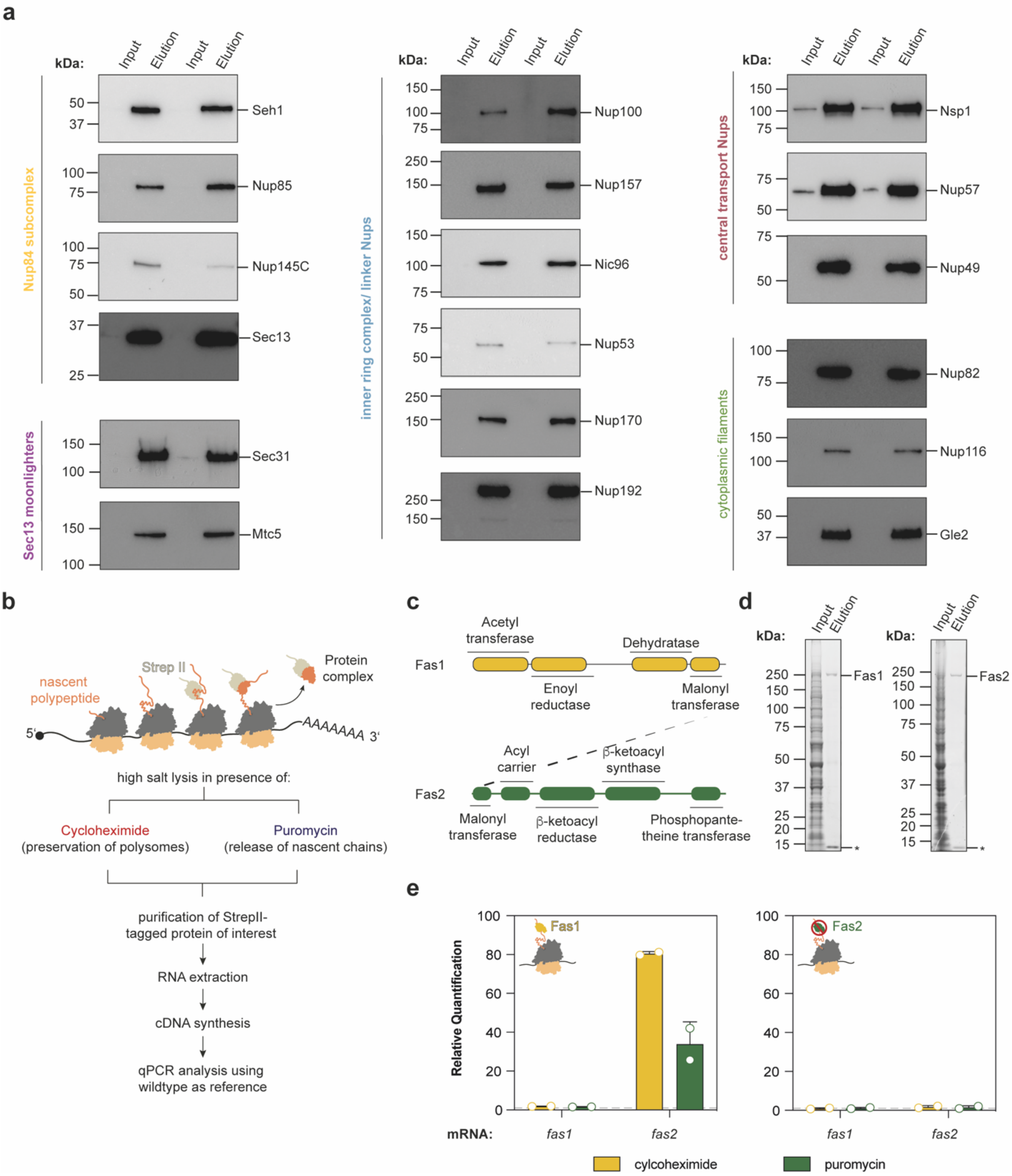


**Supplementary Figure 1: Establishment of a RIP-qPCR pipeline for Nups. a,** Representative Western Blots of the bait purifications. Proteins were detected using a primary anti-Strep antibody coupled to a secondary anti-rabbit antibody conjugated to horse radish peroxidase (HRP). **b,** Work flow of our RIP-qPCR experiment. **c,** Domain diagram of Fas1 and Fas2. The C-terminal malonyl transferase domain of Fas1 engages co-translationally with the N-terminal malonyl transferase domain of Fas2. **d,** Coomassie-stained SDS-PAGE of Fas1 and Fas2 that were affinity purified from crude lysate. Streptactin contamination is indicated with an asterisk (*). **e,** RIP-qPCR experiment of Fas1 and Fas2 showing that Fas1 could enrich for *fas2*-mRNA in a puromycin-sensitive manner. Bar plots show mean ± SD of the two biologically independent replicates of Fas1-IP and Fas2-IP, respectively.

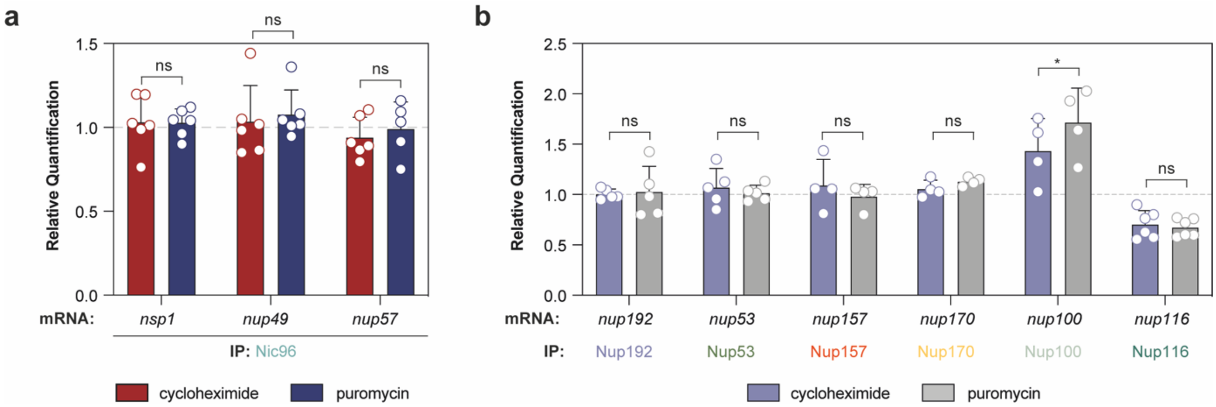


**Supplementary Figure 2: Extension of RIP-qPCR data. a,** RIP-qPCR with affinity purified Nic96 does not enrich mRNAs encoding any components of the CTN. **b,** RIP-qPCR experiments for the indicated Nups do not or only weakly enrich for their own mRNA. Bar plots show mean ± SD of four to six biologically independent experiments. ns p>0.05,* p<0.05, ** p<0.01 (Two-sided, paired t-test).

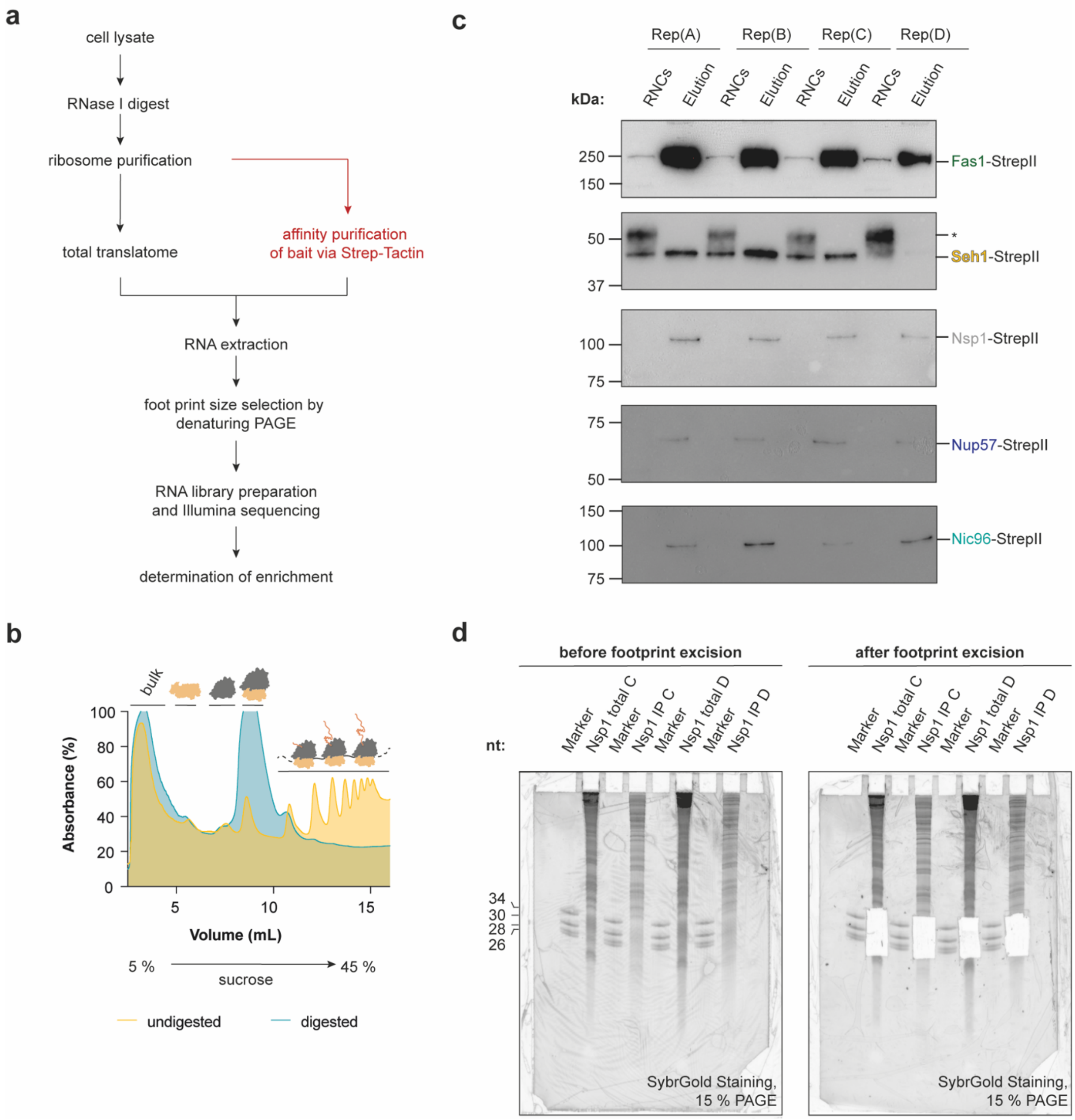


**Supplementary Figure 3: Preparation of selective ribosome profiling experiment. a,** Schematic illustration of a selective ribosome profiling experiment. **b,** Representative polysome profiling experiment of RNase I treated cell lysates. While, undigested cell lysate contains polysomes, 20 U/A of RNase I converts the majority of polysomes into monosomes (80S ribosomes) and an RNase I resistant disome peak. **c,** Western Blot analysis of the Fas1 and tested Nups shows enrichment after affinity purification in comparison to the input consisting of ribosome-nascent chain complexes (RNCs). Proteins were detected by an anti-StrepII antibody. Asterisk (*) marks an unspecific band. Membranes show the four biologically independent replicates for the SeRP experiment. **d,** Representative denaturing PAGE gel as used for the size selection of the ribosome protected footprints by gel excision. A marker with synthetized 26-, 28-, 30- and 34-mers was loaded as reference. RNA was visualized by SybrGold.

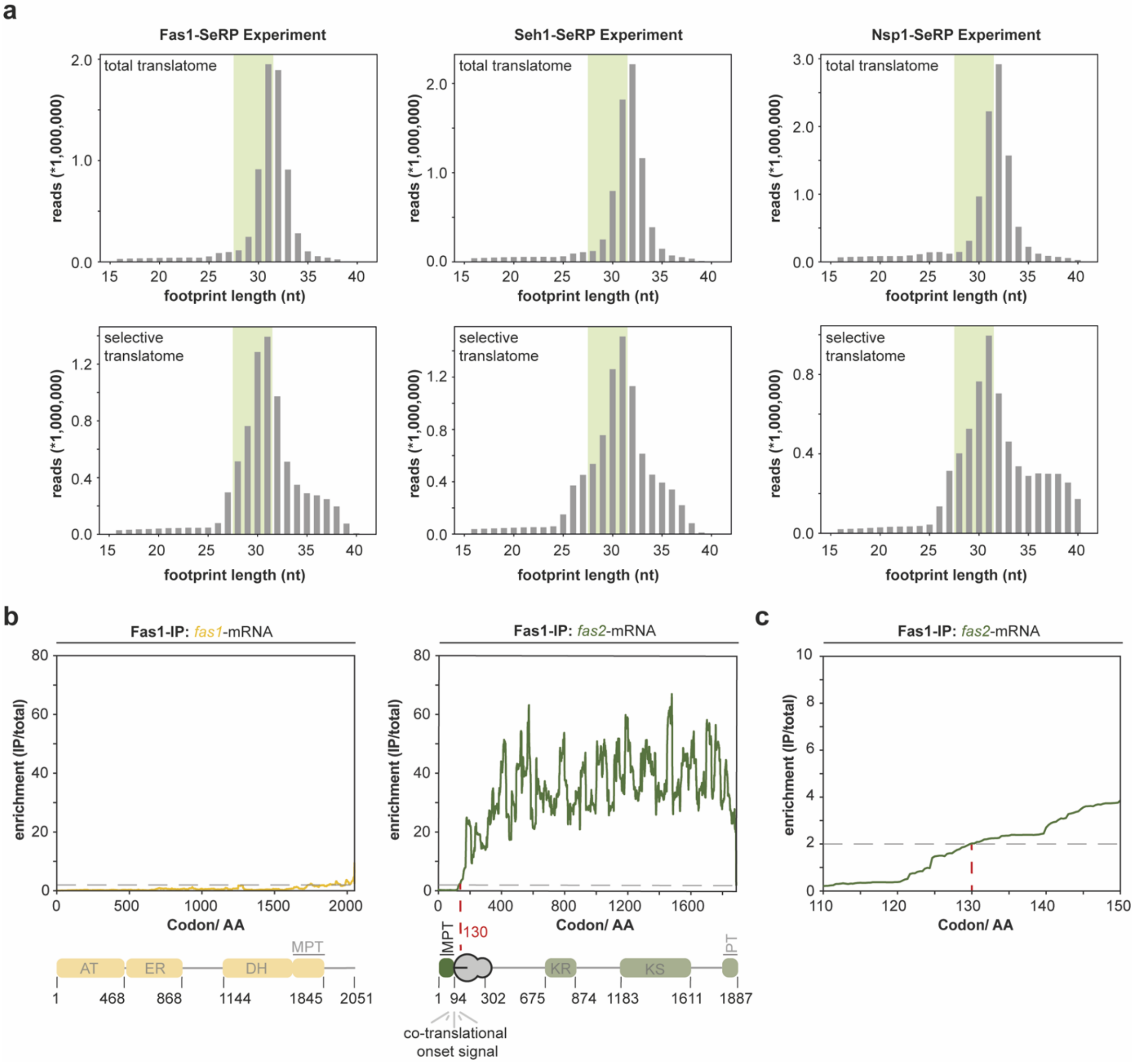


**Supplementary Figure 4: Fas1-SeRP experiments were used to benchmark the analysis pipeline. a**, Ribosome footprints of adequate length (reflecting the protected fragments) are retrieved from sequencing data. Representative footprints for Fas1-, Seh1- and Nsp1-SeRP experiments are shown. Note that footprints recovered from IPs for respective baits were derived from the total translatome. The green area (26-32 nt) highlights the footprints used for further processing. Each size distribution plot shows one of the four biological replicates. **b**, and **c**, Positive control recapitulates the co-translational interactions of Fas1 with the nascent chain of Fas2 with an interaction onset after 130 aa. Graph was generated from four biologically independent replicates. SeRP: Selective ribosome profiling, IP: immunoprecipitation, AA: amino acids.

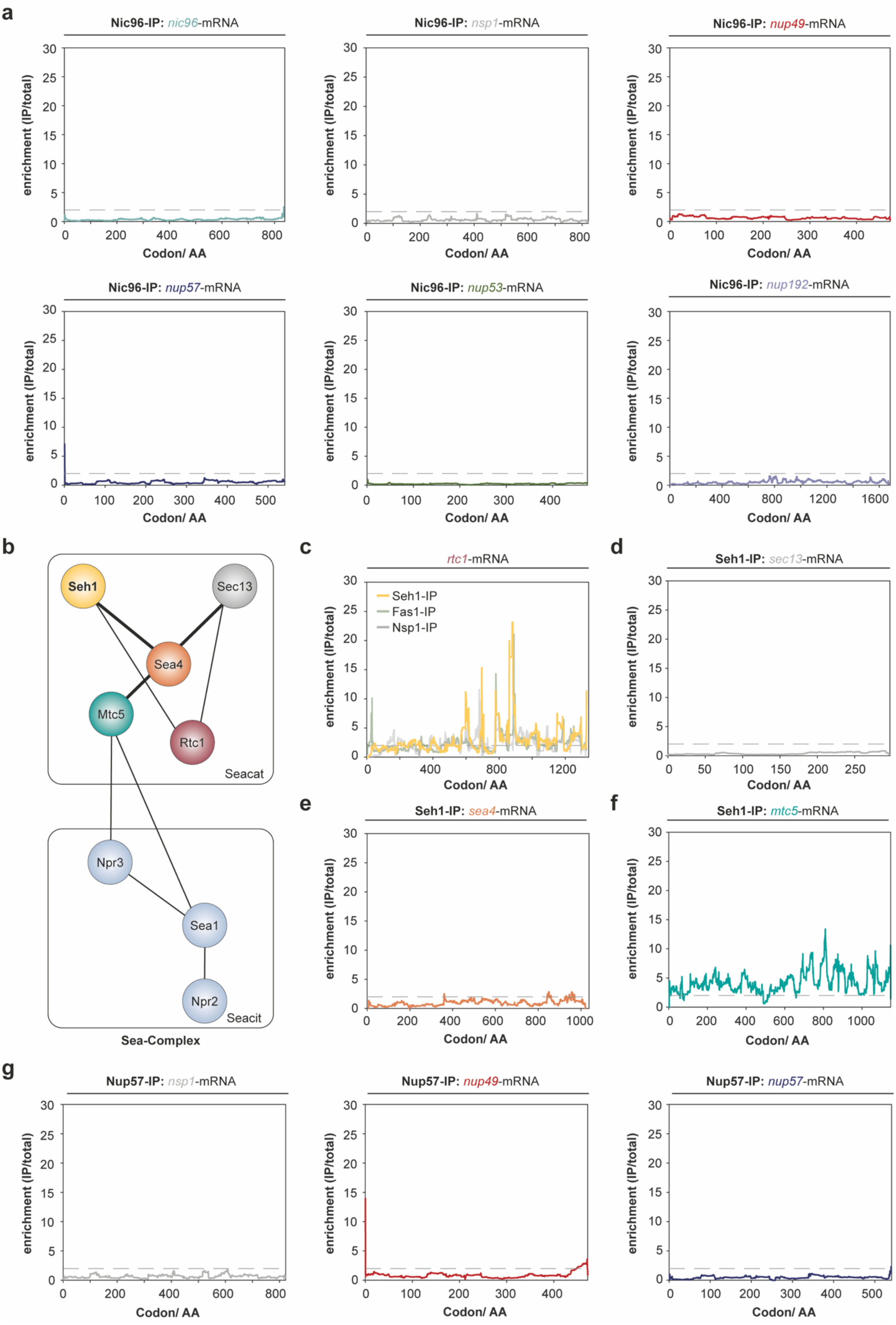
a
Nic96-IP: nic96-mRNA
Nic96-IP: nsp1-mRNA
Nic96-IP: nup49-mRNA
Nic96-IP: nup57-mRNA
Nic96-IP: nup53-mRNA
Nic96-IP: nup192-mRNA
enrichment (IP/total)
Codon/ AA
b
Seh1
Sec13
Sea4
Mtc5
Rtc1
Seacat
Npr3
Sea1
Npr2
Seacit
Sea-Complex
c
rtc1-mRNA
Seh1-IP
Fas1-IP
Nsp1-IP
d
Seh1-IP: sec13-mRNA
e
Seh1-IP: sea4-mRNA
f
Seh1-IP: mtc5-mRNA
g
Nup57-IP: nsp1-mRNA
Nup57-IP: nup49-mRNA
Nup57-IP: nup57-mRNA

**Supplementary Figure 5: Nic96, Seh1 and Nup57 do not co-translationally associate with any other interaction partner. a,** SeRP from Nic96 affinity purifications does not detect possible nascent chain interactions within any of the shown ORFs. **b,** Scheme of the Sea complex in *S. cerevisiae*. Lines connecting proteins represent physical interactions previously identified by crosslinking mass spectrometry (Algret et al., 2014). **c-f**, SeRP data within the Sea complex. **c,** Overlay of SeRP footprints within the *rtc1*-ORF from affinity purifications of Fas1, Nsp1 and Seh1. Although an enrichment over the total translatome is apparent, it is not specific for Seh1. **d-f**, SeRP experiments with affinity purified Seh1. Footprints within the *sec13-*, *sea4-*, and *mtc5*-ORFs are shown. In **f**, Analysis of the *mtc5*-footprints shows elevated signal but no clear onset, which is plausible given that Mtc5 and Seh1 do not physically interact (see **Figure S5b**). **g,** SeRP with affinity purified Nup57 does not detect any enrichment of ribosome footprints of other CTN components. The shown SeRP data was derived from four biologically independent replicates. IP: immunoprecipitation, AA: amino acids.

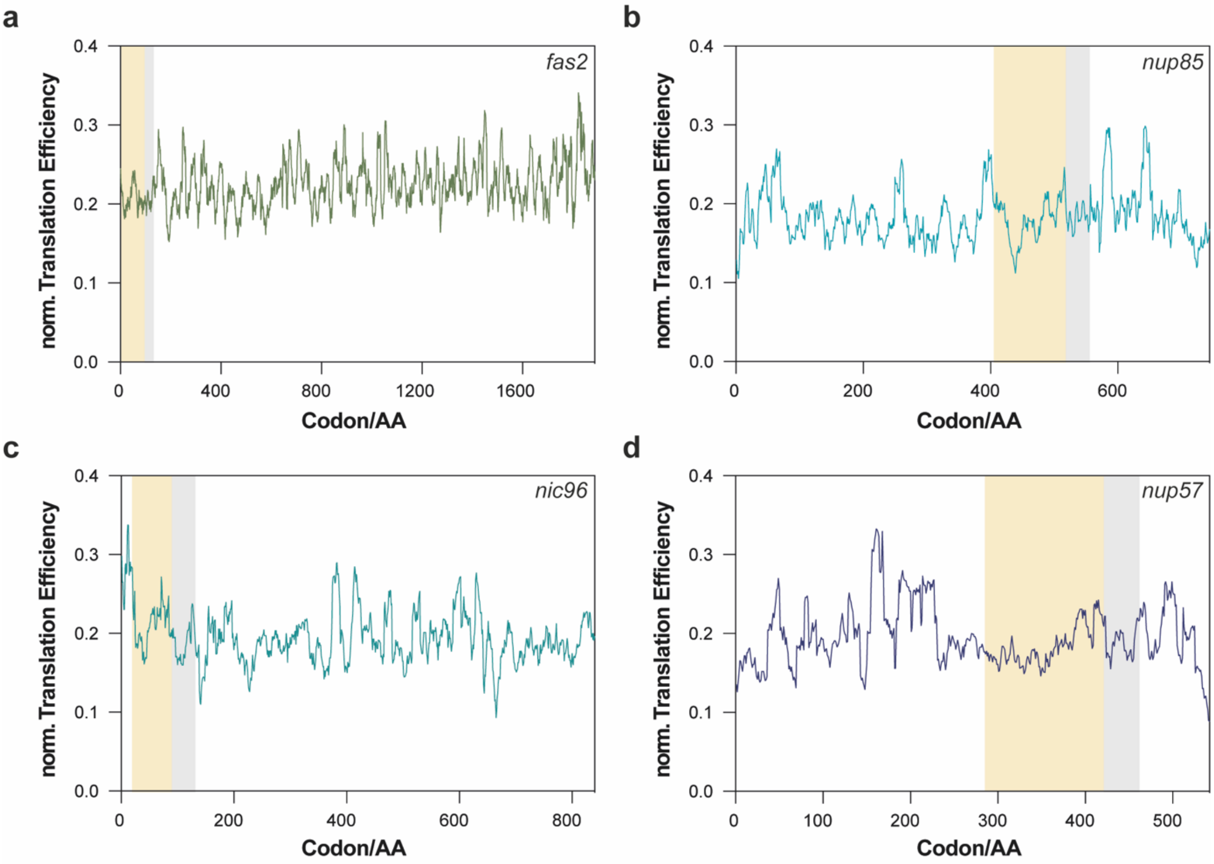


**Supplementary Figure 6:** Translation efficiency profiles for **a,** *fas2*, **b,** *nup85*, **c,** *nic96* and **d,** *nup57* (Pechmann and Frydman, 2013). The yellow area indicates the assembly domains and grey area indicates nascent chain in the exit tunnel prior to onset. Co-translational events coincide with a local reduction in translation efficiency prior to onset (grey area). norm.: normalized.

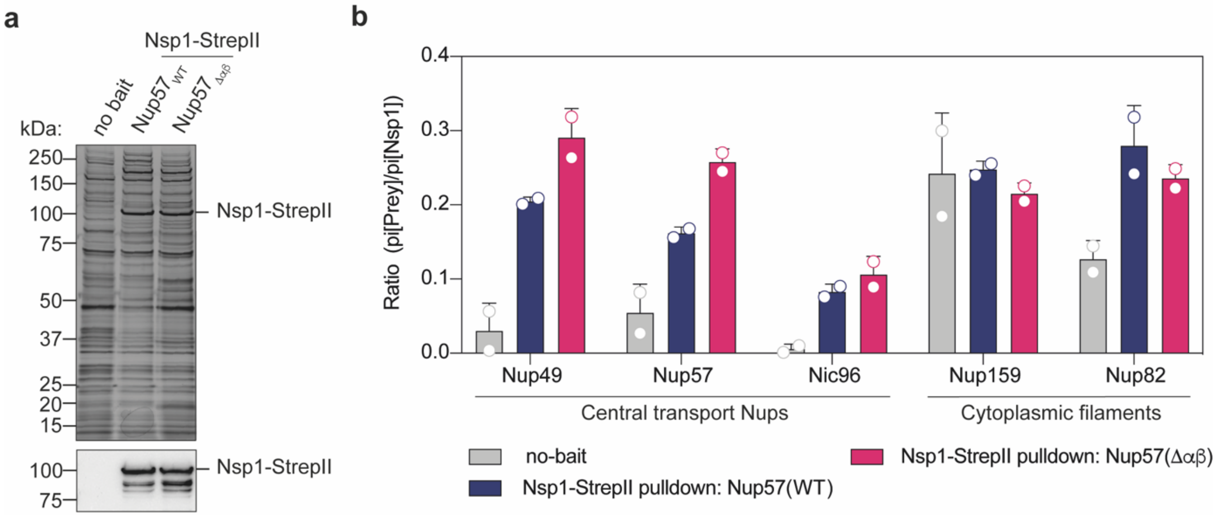


**Supplementary Figure 7: Extended data for Nsp1 pull downs from wildtype and Nup57(Δαβ)-mutant a,** Representative silver stained gel of Nsp1-StrepII pull downs (top) and Western Blot analysis using anti-StrepII antibody (bottom) revealing an enrichment of Nsp1. **b,** Mean peptide intensity of respective proteins normalized to Nsp1. Nup49 and Nup57 show increased abundances upon deletion of the Δαβ-domain of Nup57. Nic96, Nup82 and Nup159 are largely unaffected. Bar graph shows mean ± SD which was derived from two independent biologically replicates. pi: peptide intensity.
